# 3R-Compliant Murine *Ex Vivo* Platform for Scalable Brain Cancer Modeling and Drug Screening

**DOI:** 10.1101/2025.03.03.641148

**Authors:** Anna Wolfram, Vanessa Arnold, Anastassiya Moskalchuk, Carlos Romero-Nieto, Lisa Sevenich

## Abstract

Translating *in vitro* findings into *in vivo* applications remains a major challenge in brain cancer drug discovery, largely due to inadequate models for assessing neurotoxicity and tumor microenvironment (TME) dynamics. Here, we present a novel, highly modular murine *ex vivo* platform based on adult neural stem cells (aNSCs), designed for scalable, medium-throughput drug screening and mechanistic studies for brain cancer research. This system integrates progressive 2D and 3D models, including assembled models for tumor migration, cancer core models mimicking tumor localization within neural tissue and advanced cancer core models incorporating immune cells to recapitulate key TME interactions. With high reproducibility, defined cell composition and rapid formation within 72 hours, these assembloids provide a translationally relevant and ethical alternative to animal models. Aligned with 3R principles, this platform bridges the gap between *in vitro* and *in vivo* studies, offering a powerful tool for innovating brain cancer drug discovery.

## Introduction

The treatment of primary and secondary brain cancers remains challenging, with median survival averaging 10 months despite advanced multimodal regimens.^1-5^ This emphasizes the urgent need for improved treatment strategies. However, drug development faces a low success rate of only 3-7%,^6-8^ posing significant burdens in terms of time, cost, human resources and animal welfare.^9,10^ A key bottleneck is in the disconnect between *in vitro* and *in vivo* models, particularly in the assessment of neurotoxicity and tumor microenvironment (TME) dynamics.^11-13^ Rational drug design – an iterative process involving candidate synthesis, *in vitro* screening and testing in more complex models such as spheroids, organoids and animal models (**Fig. 1a**)^14^ – is hindered by poor translatability of *in vitro* results to *in vivo* systems.^15^ Many current *in vitro* models remain inadequate for assessing neurotoxicity and the complex interactions within the TME.^16-18^ In addition, initial *in vivo* drug screenings involving extensive compound libraries are increasingly constrained by ethical considerations and rising costs.^19,20^ Since 2010, the European Union’s Directive 2010/63/EU has mandated the reduction, replacement and refinement (3Rs) of animal use in scientific research.^21^ In line, significant efforts have focused on developing 3R-compliant models,^22-25^ such as *in vitro* spheroids^26-29^ (3D cancer cell aggregates) and *ex vivo* organoids^30-32^ (3D clusters of organ-specific cells) derived from animal and human tissues. These models, particularly cerebral organoids,^33-36^ have been instrumental in replicating aspects of brain development and pathology by mimicking *in vivo*-like 3D environments. However, typically cultivated from induced pluripotent stem cells (iPSCs) or embryonic stem cells (eSCs), their production is resource intensive and prone to batch variability in cell composition, limiting their utility for systematic drug discovery.^37-41^ Furthermore, most preclinical *ex vivo* models are human-based,^42-47^ while regulatory bodies including the U.S. Food and Drug Administration (FDA) and European Medicines Agency (EMA) still require animal testing for drug approval. Notwithstanding, *ex vivo* and rodent *in vivo* models can yield conflicting results due to differences in model systems (*ex vivo* vs. *in vivo*) and species (human vs. rodent).^48,49^

**Fig. 1.**
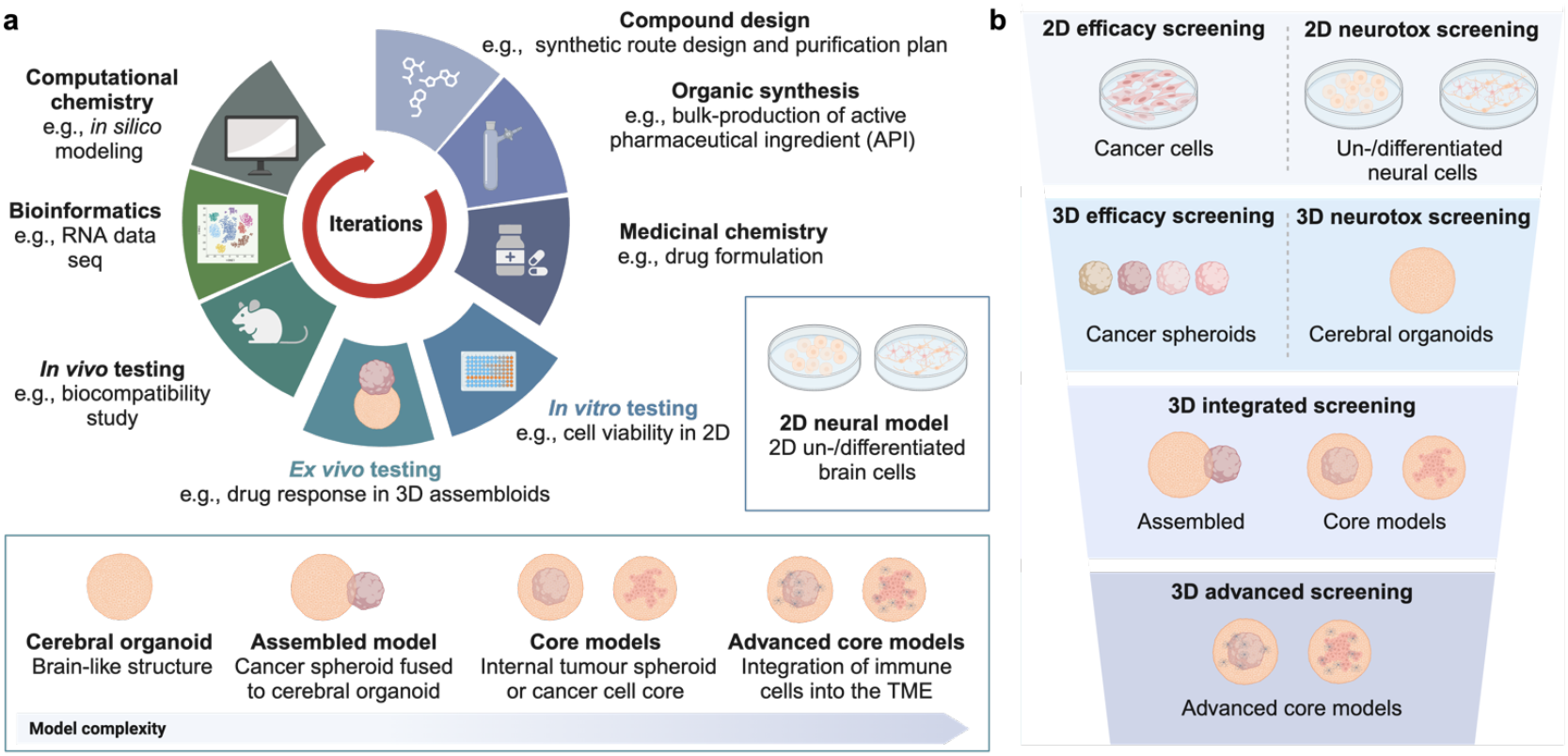
Advancing drug discovery through 3R-compliant aNSC-based models to enhance *in vitro* to *in vivo* translation. **a**, Schematic of an optimized, iterative hit-to-lead drug discovery process integrating 2D *in vitro* neural models, 3D *ex vivo* cerebral organoids and cancer-cell-integrated assembloids. These models enable neurotoxicity assessments alongside efficacy screening to improve predictive power and reduce the *in vitro* to *in vivo* translation gap. **b**, Overview of consecutive drug screening platforms with increasing biological complexity. Initial 2D neural models support high-throughput neurotoxicity screening, followed by 3D cerebral organoids for medium-throughput neurotoxicity studies. Cancer-cell assembloids enable combined efficacy and neurotoxicity assessments, while *advanced cancer core* models incorporate immune cells to account for the impact of immune-tumor dynamics in drug screening. Created with BioRender.com.

To address these limitations, we developed a 3R-compliant *in vitro* and *ex vivo* murine model based on adult neural stem cells (aNSCs)^50^ to improve cross-species compatibility and bridge the gap between human ex vivo and rodent in vivo models. As an advantage, unlike iPSCs,^51-53^ aNSCs bypass resource-intensive reprogramming and exhibit reduced batch variability, making them well-suited for systematic, reproducible drug screening. Our model constitutes a consecutive system of increasing biological complexity, ranging from 2D neural cell cultures for high-throughput neurotoxicity assays to 3D assembloids for advanced studies on tumor biology, drug efficacy and mode of action (MoA) (**Fig. 1b**). To better recapitulate TME dynamics, our murine assembloids integrate primary and secondary brain cancer cells in three distinct formats: (1) assembled models, in which pre-grown cancer spheroids fuse with cerebral organoids to study invasion and migration; (2) cancer core models, embedding single cancer cells within a cerebral organoid ‘shell’ to mimic clinically relevant tumor growth patterns; and (3) advanced cancer core models, incorporating immune cells such as microglia to simulate the immune compartment. These assembloids represent a novel murine *ex vivo* system, providing a reproducible platform to integrate cancer cells within healthy neural tissue while maintaining defined cell populations and structured tissue architectures. Among these, the *cancer core* and *advanced cancer core* models are the first to fully embed cancer cells within healthy neural tissue, replicating tumor microenvironments with defined cellular organization. Key advantages of this system include high reproducibility, defined cell composition and rapid formation, making it a powerful tool for translational research. The assembloids develop within 72 hours, enabling medium-throughput drug screening while maintaining biological complexity and scalability. By bridging human *ex vivo* and rodent *in vivo* systems, they enable preclinical drug screening and mechanistic studies. In line with the European 3R directive, these models offer an ethical and translationally relevant platform for brain cancer research, poised to innovate *in vitro* to *in vivo* translation.

## Results

### Generation and characterization of mouse adult neural stem cell culture

To identify neurogenic regions within the mouse brain, we harvested brains from C57BL/6 young adult mice aged 9-20 weeks under the internal animal tissue collection reference number at the Institute for Tumor Biology and Experimental Therapy, Georg-Speyer-Haus, Frankfurt, Germany. After fixation, we sectioned the brains into 300 μm thick slices using a vibratome. Subsequently, the sections were then stained with stem-cell markers Nestin and SOX2, neural marker NeuN and nuclear stain DAPI. We identified enriched mouse neural stem cell colonization in the lateral ventricle walls of the subventricular zone (SVZ) and the dentate gyrus (DG) of the subgranular zone (**Fig. 2a**). Adapting a previously published protocol^54,55^ (**Supplementary Method 2.2.1**), we isolated respective areas and processed them by mechanical dissection and enzymatic digestion using trypsin, followed by trypsin inhibition and filtration to collect a single-cell suspension. This suspension was then cultivated in neural stem cell basal media (**Supplementary Method 2.2.2**). Over one to two weeks, cellular debris diminished and well-shaped neurosphere cultures emerged. By day 14, these proliferating cell aggregations reached diameters of 50-100 μm and primarily consisted of stem cells and progenitor cells (**Fig. 2b**).

**Fig. 2.**
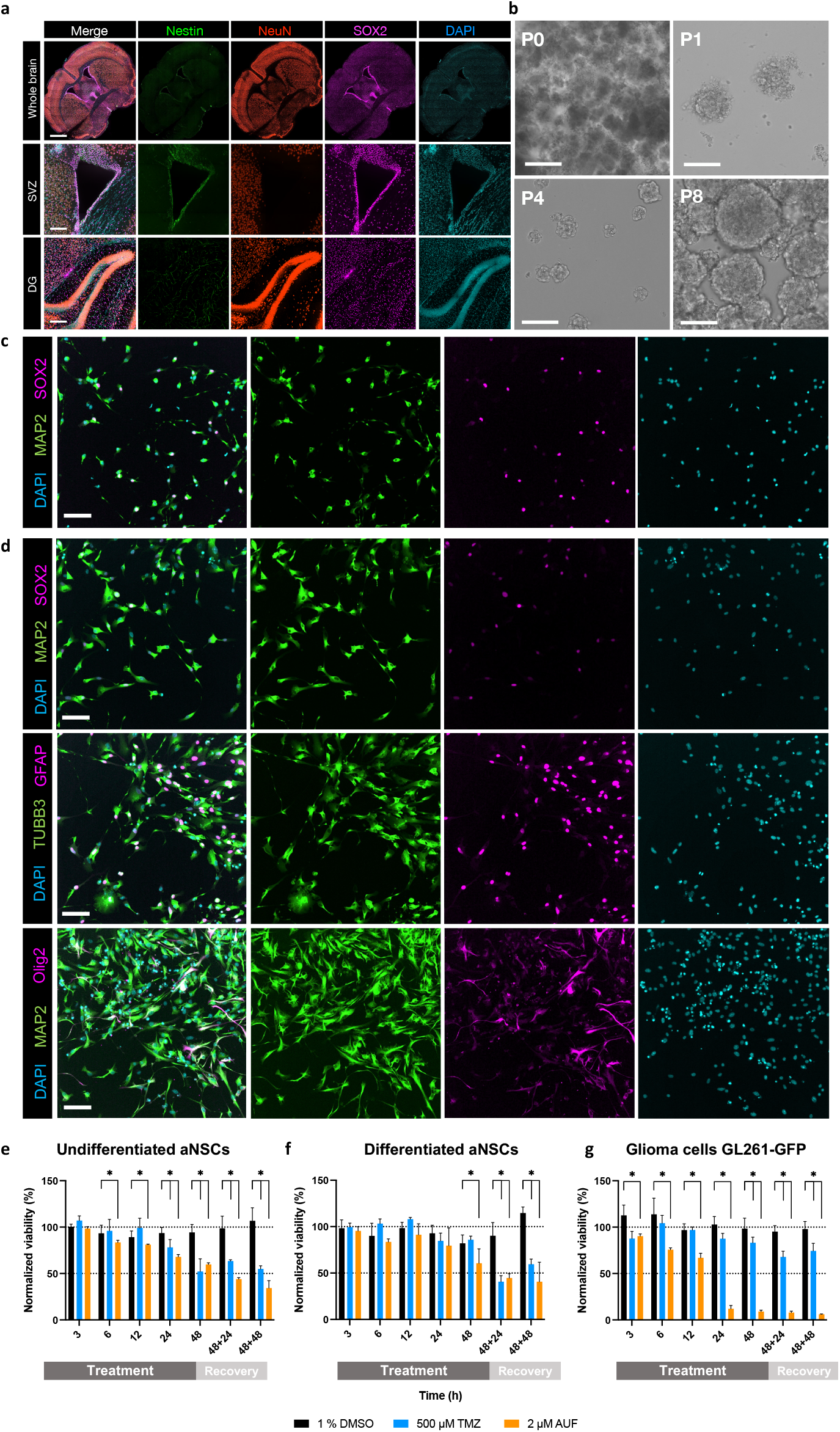
Neurosphere-derived 2D neural models. **a**, IF staining using neural stem cell markers SOX2 and Nestin and neural differentiation marker NeuN to identify neural stem cells and mature neurons in SVZ and DG. Scale bars: 500 μm and 200 μm for lower and higher magnifications, respectively). **b**, Brightfield images of aNSCs-derived neurospheres. Scale bars: 200 μm (P0) and 100 μm (P1-8), notation “P” indicates passage number). **c** and **d**, IF staining for undifferentiated neural stem cells after 24 hours (**c**) and differentiated aNSCs after 5 days (**d**) targeting the neural stem cell marker SOX2, neuronal marker MAP2, astrocytic marker GFAP and Olig2 for oligodendrocyte precursor cells (OPCs) and oligodendrocytes. Scale bar: 100 μm. **e-g**, 2D high-throughput neurotoxicity and efficacy assay. Undifferentiated (**e**) and differentiated (**f**) aNSCs, as well as GL261 glioma cells (**g**), were treated with Temozolomide (TMZ) (500 μM, blue) or Auranofin^®^ (AUF) (2 μM, orange) for 48 hours, followed by a 48-hour recovery phase. n_bio_ > 3. Controls included media-only (not shown) and vehicle-only (1% DMSO; black bars in **f-h**). Data were normalized to the media-only control at each time point (100%). Multiple t-tests compared vehicle-only to treatment conditions. Asterisks (*) indicate significance (q < 1%) after FDR correction.

To assess aNSCs-based neurospheres for stem cell characteristics and differentiation potential, we seeded 5,000 cells per well into poly-L-ornithine and laminin-coated 96-well plates overnight, according to a previously published protocol.^54^ To assess stem cell characteristics, cells were fixed and stained for SOX2 (stem cells) and differentiation markers NeuN, Olig2 and GFAP for neurons, oligodendrocytes and astrocytes, respectively. Confocal immune-fluorescence (IF) images of undifferentiated aNSCs showed high abundance of the stem cell marker SOX2, indicating robust stem cell characteristics (**Fig. 2c, Supplementary Method 2.2.3**).

To analyze differentiation potential, seeded cells were incubated in growth-factor-reduced media for 3 days before fixation and staining. aNSCs incubated with growth factor withdrawal exhibited a loss of SOX2 expression but showed a high abundance of Olig2 and GFAP, confirming the presence of oligodendrocytes and astrocytes respectively (**Fig. 2d, Supplementary Method 2.2.4**). Furthermore, the neural morphology exhibited a highly interconnected network indicative of neural differentiation. Therefore, aNSCs effectively generated the primary brain parenchyma cell types.

### aNSCs-based assay for 2D high-throughput neurotoxicity assessment

Next, we developed a 2D high-throughput neurotoxicity assay using aNSCs to enable drug screening in both undifferentiated and differentiated neural cells (**Supplementary Method 2.2.5**). Viability was assessed using an MTS-based assay, where the bio-reduction of MTS into violet formazan is measured by absorbance at 490 nm, reflecting metabolic activity and thus cell viability.

In a proof-of-concept experiment, 6×10^4^ aNSCs per well were seeded in pre-coated 96-well plates and cultured under two conditions: undifferentiated aNSCs (cells attached overnight; **Fig. 2e**) and differentiated aNSCs (cells differentiated for three days into neurons, astrocytes and oligodendrocytes; **Fig. 2f**). As a reference, 1×10^5^ GL261 glioma cells per well were seeded and allowed to attach overnight (**Fig. 2g, Supplementary Method 2.2.6-2.2.7**). Cells were treated with temozolomide^1,2^ (TMZ, 500 μM) – the standard-of-care (SOC) chemotherapy for glioblastoma (GBM) – or Auranofin^56,57^ (AUF, 2 μM) – an FDA-approved antirheumatic drug currently under investigation for oncology applications. Concentrations were selected based on previously published therapeutically active concentrations.^58-60^ After 48 hours of drug exposure, all conditions were replaced with fresh media to simulate a drug clearance phase (recovery) and viability was assessed at multiple time points: 3, 6, 12, 24, 48 hours (treatment) and 48+24 and 48+48 hours (recovery). All results were normalized to the media-only control at each time point (set to 100%). Controls included media-only (not shown) and vehicle-only (1% DMSO; **Fig. 2f-g**). Both TMZ and AUF exhibited comparable neurotoxicity in undifferentiated and differentiated aNSCs, with a greater impact on undifferentiated cells, consistent with the higher sensitivity of proliferative populations to chemotherapeutic agents. Neurotoxicity in differentiated cells showed a delayed onset, becoming evident at 48 h, with partial recovery at 48+48 h. In contrast, the effects of TMZ and AUF diverged in glioma cells (GL261). TMZ-treated glioma cells exhibited only a mild viability reduction (∼25%) after 48 hours and started to recover by 48+48 h, indicating limited efficacy as seen in the clinic. AUF, however, induced rapid and sustained cytotoxicity, with an 80% viability loss at 24 hours and no recovery during the post-treatment phase. This aNSC-based assay provides a simple and scalable platform for high-throughput neurotoxicity screening, allowing direct comparison of drug effects in healthy and malignant cell populations.

### aNSCs 3D differentiation to access mature cerebral organoids

Leveraging the inherent high stem cell character of aNSCs, we aimed to generate differentiated 3D cerebral organoid cultures (**Supplementary Method 2.2.8**). Utilizing a modified protocol,^54^ we seeded 10,000 aNSCs per well into ultra-low adherent (ULA) 96-well plates to promote the formation of well-defined spherical organoids. For differentiation, the 3D aggregates were incubated in media with pre-defined differentiation factors on a CO_2_-resistant orbital shaker at 37°C with 5% CO_2_. Once organoids exceeded 300 μm in diameter, they were transferred using single-use 3 mL Pasteur pipettes to 6**-**well suspension plates or the CERO 3D Incubator & Bioreactor (OMNI Life Science, short: CERO). Continued gentle swirling on the shaker or in CERO encouraged uniform growth and maturation of the organoids into well-defined spheres (**Fig. 3a**).

**Fig. 3.**
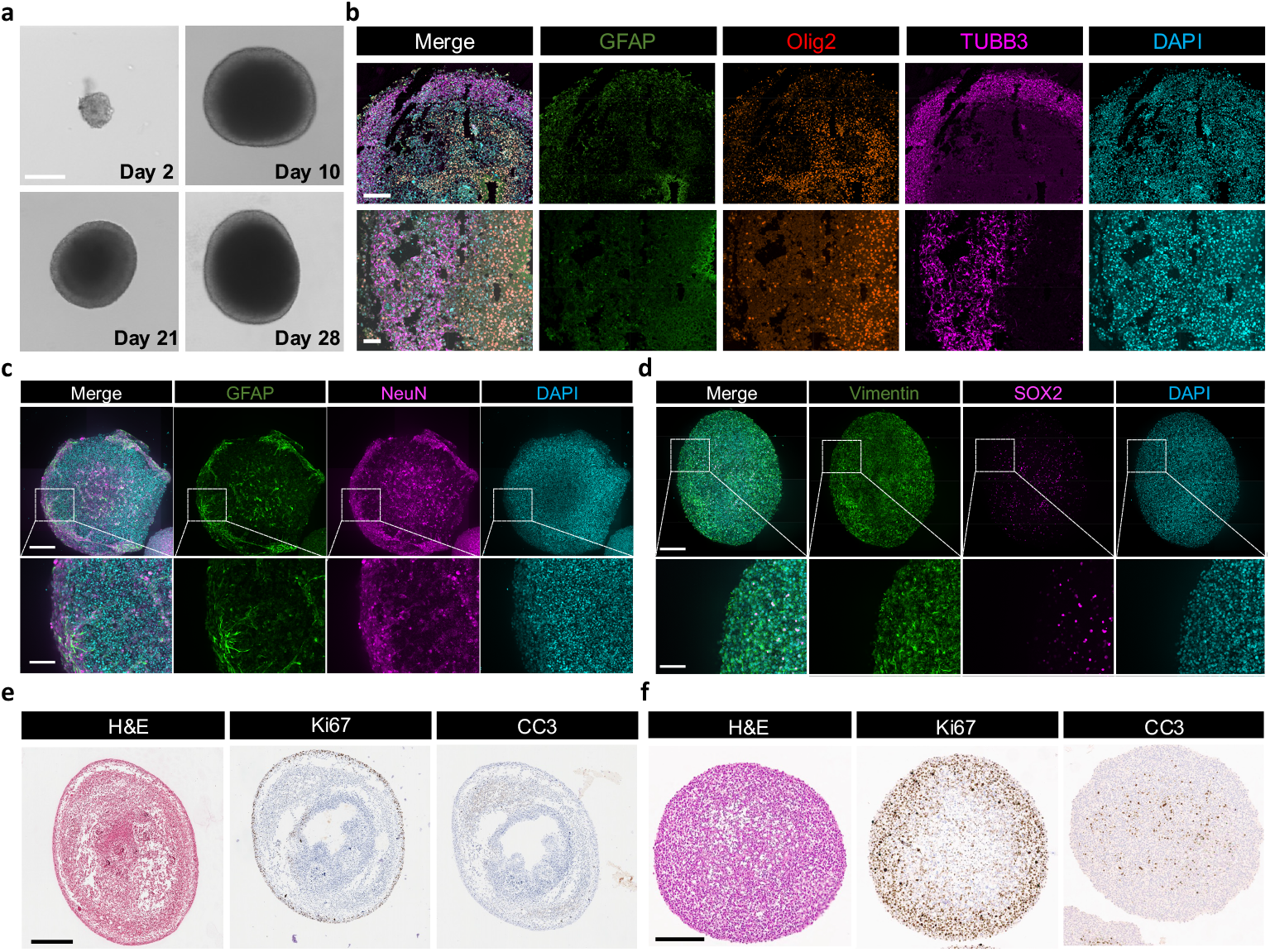
aNSC-derived large cerebral organoid culture. **a**, Growth stages of cerebral organoids derived from aNSCs at 2, 14, 21 and 28 days. Scale bar: 200 μm. **b**, Thin sections of cerebral organoids showing neural maturation markers for astrocytes (GFAP), neurons (TUBB3) and oligodendrocyte lineage cells (Olig2). Scale bar: 200 μm, 50 μm. **c and d**, IF staining of whole 8-week-old organoids showing differentiation markers GFAP and NeuN for astrocytes and neurons (**c**) and neural stem cell markers Vimentin and SOX2 (**d**). Scale bars: 225 μm, 100 μm. **e** and **f**, IHC analysis of 12-week-old organoid sections, stained for Ki67 and CC3 in orbital shaker (**e**) and CERO (**f**) cultures. Scale bars: 500 μm (**e**) and 300 μm (**f**).

### Large organoid preparation and analysis

To access large cerebral organoids up to 2 mm in diameter, we extended the culture period to three months before fixing them. We employed filter cassette-based fixation techniques optimized for organoid fixation, facilitating subsequent paraffin embedding and sectioning (**Supplementary Method 2.2.9**). Organoids underwent IF (**Fig. 3b-d**) and immunohistochemical (IHC) staining (**Fig. 3e, f**), revealing distinct morphological and cellular distribution patterns. Antibody staining confirmed the presence of major brain parenchyma cell types. Neuronal networks were well-developed, astrocytes exhibited star-like shapes and Olig2 staining identified oligodendrocyte precursor cells (OPC) and oligodendrocytes. Stem cell populations remained evident within organoids after 2 months of differentiation.

Hematoxylin and eosin (H&E) staining revealed that organoids incubated in the CERO system exhibited a uniform, highly differentiated morphology, whereas those grown in a shaker system appeared elongated and less uniform. Cleaved Caspase-3 (CC3) staining indicated minimal apoptosis in CERO organoids and antigen Kiel 67 (Ki67) staining showed high, evenly distributed proliferation, in contrast to the rim-restricted proliferation observed in shaker-grown organoids. These findings suggest that the floating rotation systems supports the generation of more homogeneous organoids with structural characteristics that more closely resemble in vivo tissue.

### 3D *assembled* models capturing tumor spheroid-cerebral organoid interactions

To model non-cancerous–cancerous tissue interactions, we developed brain cancer assembloids using two approaches: (1) *assembled* models, where cancer spheroids fuse with organoids to study cell migration and (2) *cancer core* models, where single cancer cells are embedded in cerebral organoid shells, mimicking tumor architecture.

In the *assembled* model, organoids grown to 0.8-1 mm in diameter were co-cultured with pre-formed cancer spheroids. We seeded 10,000-40,000 cancer cells (depending on the cell line) in ULA plates to form spherical 3D cultures and monitored growth with bright-field and fluorescence microscopy for cell lines expressing fluorescent markers (GFP, RFP, BFP). After 48–72 hours, cancer spheroids were combined with four-week-old cerebral organoids by transferring single organoids into fresh ULA wells with a selected cancer spheroid (**Supplementary Method 2.2.10**). To establish cell proximity, plates containing one organoid and one spheroid per well were spun down and incubated at 37°C with 5% CO_2_.

In a proof-of-concept time lapse (TL) live imaging study, we used cancer spheroids from four cell lines: GL261-GFP^61,62^ (glioma, seeded at 10,000 cells per spheroid), LLC1-GFP^63^ (Lewis lung carcinoma, 10,000 cells/ spheroid), MK1735-BFP^64^ (melanoma-to-brain, MK-BFP, 10,000 cells/ spheroid) and 99LN-BFP^65^ (breast-to-brain, seeded at 40,000 cells/spheroid). The *assembled* model in an ULA plate was placed in the CellVoyager™ CQ1 Benchtop High-Content Analysis System (Yokogawa) and incubated in a humidified chamber at 37°C with 5% CO_2_ (**Supplementary Method 2.2.11**). Tumor spheroid-cerebral organoid interactions were monitored bi-hourly over 72 hours using spinning disc confocal microscopy to visualize dynamic integration patterns.

TL imaging revealed distinct cancer cell line-dependent infiltration patterns. Specifically, glioma spheroids exhibited a large cell-cell interaction surface area and an infiltrative border region (**Fig.4a, Supplementary Video 1**). Lewis lung carcinoma cells demonstrated highly invasive behavior, with metastatic spread extending distally from the primary spheroid site (**Fig. 4a, Supplementary Video 2**). In contrast, melanoma-to-brain spheroids showed a more localized growth pattern, with cerebral cells encircling the tumor spheroid (**Fig. 4a, Supplementary Video 3**). Meanwhile, breast-to-brain cancer cells formed densely packed spheroids, with delayed tumor infiltration into cerebral tissue (**Fig. 4a, Supplementary Video 4**). To further assess 99LN cell infiltration, we extended co-culture to 120 hours before light-sheet microscopy (LSM) analysis (**Supplementary Method 2.2.12**). LSM imaging confirmed 99LN cell infiltration, with an additional 48 hours leading to increased metastatic spread, including distal tumor herds far from the primary tumor location (**Fig. 4b,c**).

**Fig. 4.**
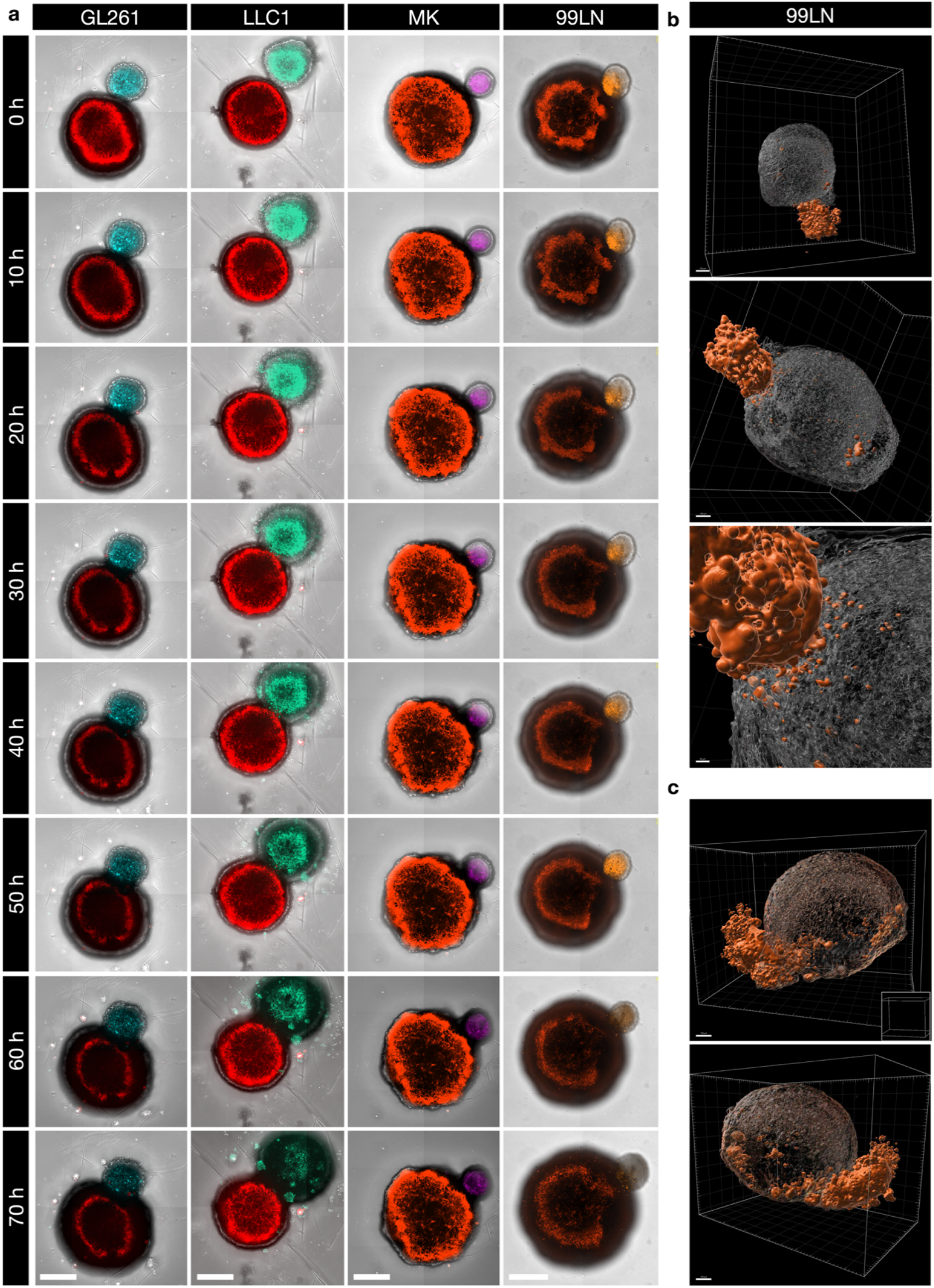
Infiltrative potential of various cancer spheroids co-cultured with aNSC-derived cerebral organoids. **a**, Time lapse imaging of cerebral organoids co-cultured with tumor spheroids derived from primary and secondary brain cancer cell lines, including glioma (GL261, blue), lung-to-brain (LLC1, turquoise), melanoma-to-brain (MK, magenta) and breast-to-brain (99LN, orange), observed over 72 hours (scale bar: 500 μm). **b** and **c**, Three-dimensional visualization of a cerebral organoid fused with a breast-to-brain (99LN, orange) spheroid, indicating cancer cell migration into the cerebral tissue over 5 days. Scale bar: 100 μm.

### Developing *cancer core* models with defined and reproducible tumor growth patterns

To enhance cell-cell interactions within assembloids, we developed an alternative approach that embeds cancerous tissue within the core of an assembloid, creating the *cancer core* model. This four-step method includes (1) generation of single tumor cell suspensions, (2) preparation of single-cell organoid suspensions, (3) tumor cell–organoid co-seeding and (4) model growth (**Supplementary Method 2.2.13**).

Single-cell suspensions were prepared from primary (e.g., GL261) and secondary cancer cell lines (e.g., 99LN, MK, LLC1) through sequential trypsinization, centrifugation and resuspension. Cerebral organoids were enzymatically dissociated, mechanically triturated, digestion inhibited, centrifuged and resuspended. As a control, cerebral organoid suspensions were seeded alone (100,000 cells per well; **Supplementary Fig. S1**), reorganizing into distinct, round organoid-like structures within 24 hours, with high viability and rapid tissue re-association. Cancer and cerebral cells were co-seeded at defined ratios (1:2 to 1:100) in ULA plates, incubated at 37°C, 5% CO_2_ on an orbital shaker and monitored via fluorescence microscopy to assess tumor architecture.

At a 100:1 ratio, cancer cells were dispersed throughout the organoid, forming a patch-like pattern (**Fig. 5, Supplementary Video 5**). A 10:1 ratio led to a dense tumour core surrounded by a cerebral shell (**Supplementary Fig. S1**), while 4:1 produced an extended tumor core covered by a cerebral layer (**Fig. 6, Supplementary Fig. S2**). At 2:1, cancer cells were interspersed throughout the assembloid, generating a mosaic-like pattern (**Supplementary Figs. S3, S4**).

**Fig. 5.**
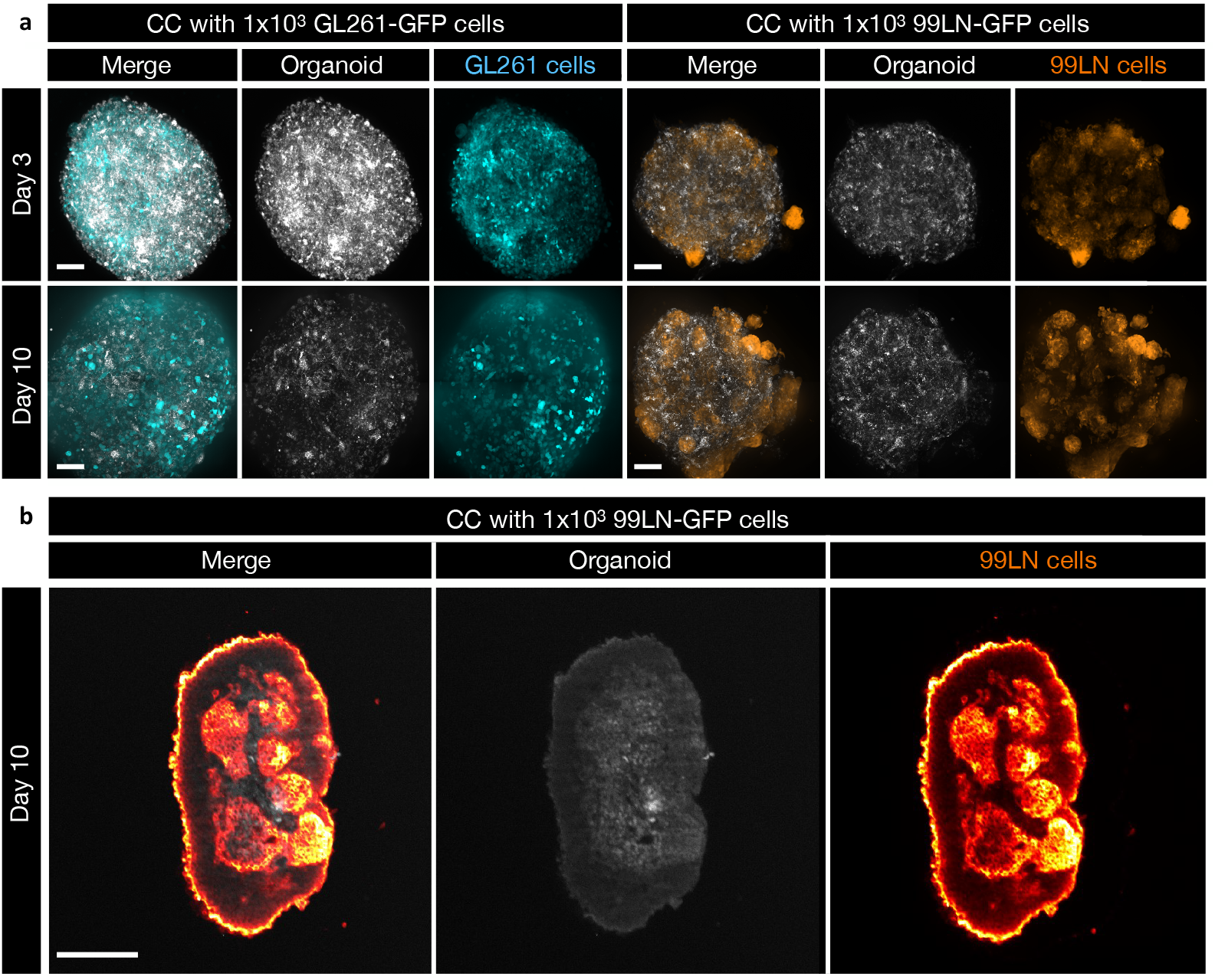
Characterization of *cancer core* models and control cerebral organoids. **a**, IF images of 1:100 *cancer core* co-cultures at 3 and 10 days show GL261-GFP (left) and 99LN-BFP (right) interactions, revealing cancer cell background-dependent growth behavior, with GL261 exhibiting highly infiltrative growth and 99LN forming lobular tumor areas (scale bar: 100 μm). **b**, 3D light-sheet imaging of a 1:100 cancer core co-culture (RFP-labeled cerebral organoid cells and 99LN breast-to-brain metastatic cancer cells) at 10 days reveals the patch-like growth of cancer cells within the organoid architecture. Scale bar: 100 μm.

**Fig. 6.**
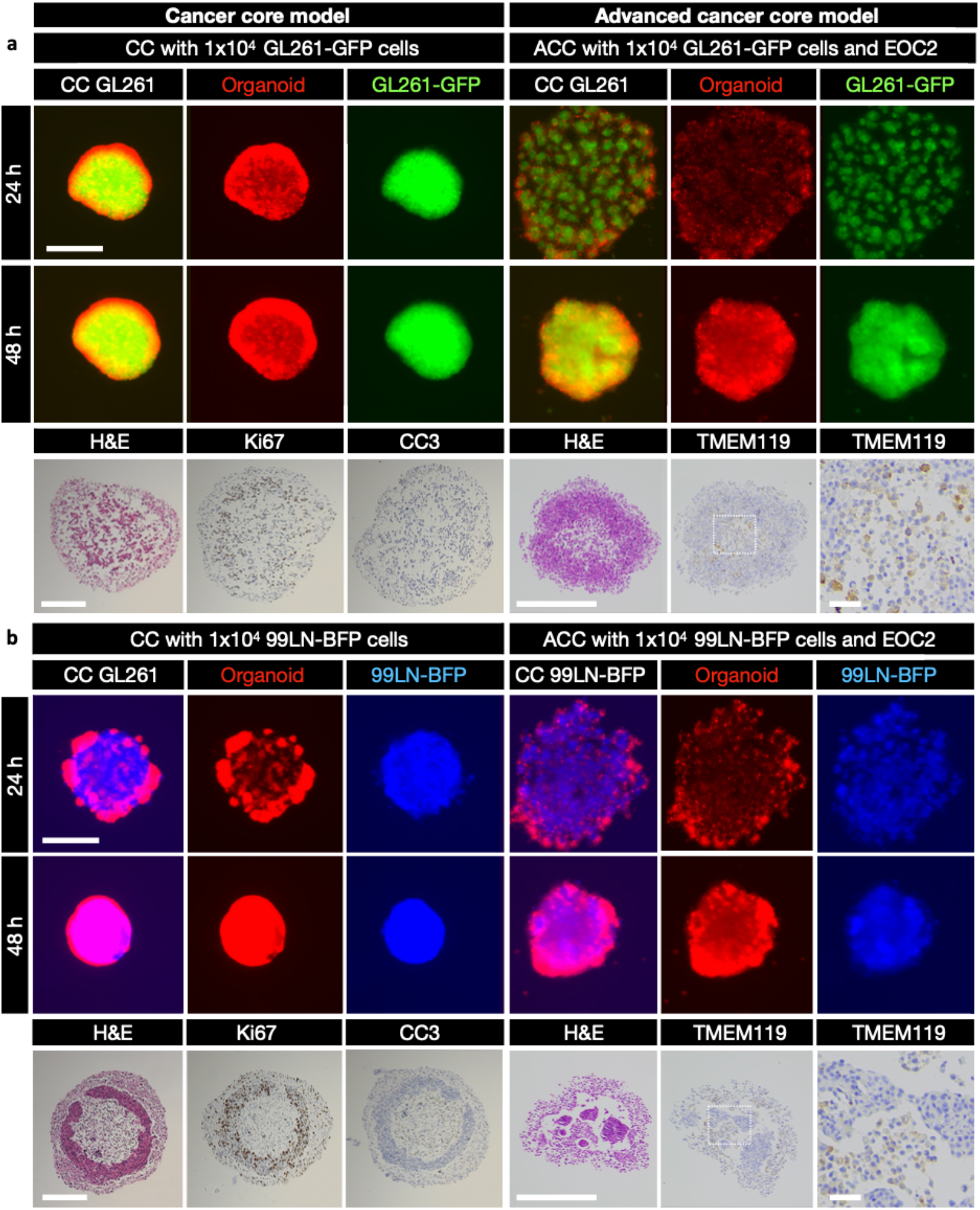
Immune cell integration drives distinct tissue architectures in glioma and breast-to-brain metastasis models. *Cancer core* models (8×10^4^ cerebral organoid cells and 2×10^4^ tumor cells; 4:1 ratio, left) were compared to *advanced cancer cores* with microglia (8×10^4^ cerebral organoid cells, 2×10^4^ tumor cells and 4×10^4^ EOC2 microglia; 4:1:2 ratio, right). **a**, GL261-GFP glioma model: RFP-labeled cerebral organoid cells (red) and GL261-GFP tumor cells (green) were visualized at 24 (top row) and 48 hours (middle row). IHC staining for HE, Ki67 and TMEM119 (bottom row) shows tumor morphology, proliferation and microglial content. Microglia are integrated into the tumor mass. **b**, 99LN-BFP breast-to-brain metastasis model: RFP-labeled cerebral organoid cells (red) and 99LN-BFP tumor cells (blue) were visualized at 24 (top row) and 48 hours (middle row). IHC staining for HE, Ki67 and TMEM119 (bottom row) indicates tumor morphology, proliferation and microglial content. Microglia are excluded from the tumor core and accumulate at the tumor rim, with evidence of microgliosis. Scale bars: 500 μm and 200 μm for TMEM119 zoom.

Cancer cell infiltration also varied by biological background, with glioma cells exhibiting diffuse invasion into neural tissue, while breast-to-brain cancer cells formed distinct clusters. LSM confirmed the herd-like growth pattern of brain-metastatic tumor cells, revealing well-defined tumor structures (**Fig. 5b**).

IHC staining of 10:1 *cancer cores* revealed highly proliferative tumor regions and cell type-specific infiltration patterns within neural tissue. Consistent with IF results, GL261 cells exhibited widespread distribution throughout the co-culture, while 99LN cells formed distinct ring-like cores. Ki67 staining confirmed high proliferation within cancerous tissue, whereas CC3 staining indicated minimal apoptosis in non-cancerous regions (**Fig. 6**).

By overcoming the limitations of traditional assembloids, particularly the minimal interface between pathological and neural tissue, the *cancer core* model more accurately recapitulates clinically observed tumor patterns, from diffuse infiltration to well-defined boundaries. This scalable and reproducible platform provides a controlled system for studying cancer progression, invasion and therapeutic responses.

### Integrating microglia to enhance tumor microenvironment modeling

To further enhance the complexity of our *cancer core* model and mimic the immune component of the TME, we incorporated microglia (EOC2) to create the *advanced cancer core* model (**Supplementary Method 2.2.14**). This process followed the established protocol for generating single-cell suspensions of cerebral organoids and cancer cells, with EOC2 microglia added after mechanical dissociation and filtration to obtain a single-cell suspension. The respective cell suspensions were seeded together in ULA 96-well plates at defined ratios of non-cancerous brain cells, cancer cells and EOC2 microglia (2:1:1, 4:1:1 and 4:1:2). Control co-cultures, without microglia, were prepared in parallel for comparison (**Fig. 6**).

To support microglia survival and activity, co-cultures containing EOC2 were maintained in a specialized medium supplemented with TGF-β and interleukin-34 (IL-34). The plates were incubated at 37°C and 5% CO_2_ on an orbital shaker, with fluorescence microscopy used to monitor the cultures over time. Models containing EOC2 displayed differences in cell distribution at 24 hours, with a mosaic-like pattern observed in *advanced cancer cores*; however, this effect reduced by 48 hours (**Fig. 6, Supplementary Fig. S2, S3, S4**).

80 hours post-seeding, the co-cultures were fixed and processed for IHC. The addition of microglia resulted in a more complex TME. IHC staining with anti-TMEM119 revealed EOC2 microglia integrated within the infiltrative tumor mass of the GL261-GFP glioma models, reflecting high microglia infiltration as seen in patient samples (**Fig. 6a**). In contrast, microglial exclusion was observed in the advanced 99LN-BFP breast-to-brain metastasis models, consistent with the distinct immune landscapes of these tumor types (**Fig. 6b**). These findings demonstrate the potential of the *advanced cancer core* model to closely mimic clinical histology, making it highly suitable for applications such as tumor microenvironment-targeted therapy screening and multimodal treatment evaluation.

This development enables more adaptable and customizable models, providing a valuable platform for rational drug screening strategies. These improvements support the exploration of a range of therapeutic interventions tailored to the specific characteristics of the TME.

### Robust FACS-based strategy for analyzing cell-specific drug responses in *cancer core* models

In a proof-of-concept experiment, we developed a robust workflow to isolate and quantify healthy and pathological cell populations following a 96-hour co-culture period. We established a fluorescence-activated cell sorting (FACS)-based strategy to separate viable cell populations in glioma core models (**Supplementary Fig. S5**). Up to 10 glioma cores were pooled, centrifuged, enzymatically dissociated with trypsin, neutralized with trypsin inhibitor, resuspended in PBS and immediately subjected to cell sorting using a BD FACSAria™ III Cell Sorter (**Supplementary Method 2.2.15**). Our gating strategy used wild-type cerebral organoids prior to DAPI staining to define unstained healthy populations, while RFP-positive cerebral organoids stained with DAPI enabled dead cell exclusion and visualization of viable RFP-positive neural cells. GL261-GFP glioma spheroids were stained with DAPI to exclude nonviable cells. Gating parameters enabled precise separation of viable DAPI-negative, RFP-positive neural cells from DAPI-negative, GFP-positive glioma cells (**Supplementary Fig. S5**). The same approach was successfully applied to breast-to-brain core models (**Supplementary Fig. S6**), demonstrating the robustness of this method for cell-type-specific separation.

Following validation of the separation strategy, we generated new glioma core models, allowing for a 72-hour co-culture period prior to a 24-hour treatment with conditions of interest: vehicle control (1% DMSO v/v), 2 μM AUF and 5 μM AUF. Drug concentrations were selected based on prior 3D tumor spheroid screenings and each condition was tested in replicates to minimize variability. Model integrity was evaluated using benchtop fluorescence microscopy (**Supplementary Fig. S7**), vehicle control-treated models exhibited well-defined, rounded edges, whereas AUF-treated cores showed compromised rim integrity, suggesting drug-induced structural disruption. Subsequently, control and AUF-treated glioma cores underwent FACS-based cell population separation. Isolated populations can be used for further downstream analyses, including RNA sequencing, to investigate cell-type-specific drug responses and mechanisms of action.

In summary, this workflow provides a robust method for isolating and analyzing cell-type-specific drug responses in complex *cancer core* models, enabling downstream molecular characterization and mechanistic insights into treatment effects.

## Discussion

In this study, we developed a suite of 3R-compliant assembloid models – *assembled, cancer core* and *advanced cancer core* – to overcome key limitations in preclinical brain cancer research.

Conventional models often fail to replicate critical features of the brain TME, such as cellular heterogeneity, tumor-immune interactions and tumor infiltration, while also lacking robust methods to assess drug-induced neurotoxicity. Translational gaps between rodent *in vivo* and human *ex vivo* systems further hinder the development of clinically relevant therapies, highlighting the need for platforms that bridge species and model-specific differences. Our *ex vivo* models address these challenges by providing scalable systems that replicate key features of the brain TME and enable modular remodeling of its complexity. The *assembled* model supports studies of cancer cell infiltration, the *cancer core* facilitates studies of tumor growth within a full cerebral ‘shell’ and the *advanced cancer* core further incorporates immune cells, such as microglia, replicating clinically observed features, such as microglial exclusion in metastatic niches and increased microglial infiltration in GBM. By bridging translational gaps between mouse *in vivo* and human *ex vivo* systems, they enable direct comparisons across species and models. With high reproducibility, rapid generation times (72 hours) and scalability for medium-throughput screening, these models address both scientific and logistical challenges, offering significant improvements over surface-based assembloid systems. Beyond drug screening, these models support multi-omic analyses of defined cell populations, including transcriptomics, proteomics, metabolomics and lipidomics, enabling detailed evaluations of drug efficacy, neurotoxicity and mode of action in both healthy and pathological brain tissues.

Despite their strengths, the increased biological complexity in *advanced cancer cores* necessitates rigorous validation and standardization, particularly when scaled for medium-throughput applications. Further studies are needed to refine their translational potential and validate their performance across diverse experimental conditions.

In the future, we plan to expand the *advanced cancer core* models to include other immune cell types, such as T cells, which will allow for comprehensive studies of tumor-immune interactions and facilitate evaluations of combination therapies, including immunotherapies. Additionally, we aim to translate the core models into human iPSC-derived systems to develop patient-specific platforms with enhanced clinical relevance. Incorporating human-derived cerebral organoids, immune components and patient-specific tumor tissue will enable personalized medicine applications and provide a more accurate replication of human disease pathology.

In conclusion, unlike previously published approaches, this study presents the first methodology for generating assembloids with fully embedded cancer cells, marking a significant advancement. These assembloids further integrate tumor-immune interactions, recapitulate clinically relevant architectures and enable modular remodeling of tissue complexity. Their rapid formation (72 hours), high reproducibility and scalability support medium-throughput drug screening while preserving critical tumor microenvironment features. Their modular design allows for broader applications in other solid cancers by exploiting tissue-specific stem cells, thereby potentially expanding their utility beyond brain cancer research. By bridging translational gaps between *in vivo* and *ex vivo* models, these assembloids provide a more physiologically relevant platform for studying tumor biology, drug responses and immunotherapy. With their potential to accelerate therapeutic discovery, improve translational accuracy and potential future application in personalized medicine, these models are set to innovate the preclinical pipeline and address critical questions in oncology and beyond.

## Acknowledgment

We thank Petra Dinse (Georg-Speyer-Haus) and Jenny Hetzer (M3 Research Center) for their expertise and support in histology. We are grateful to Nicola Bell for her contributions during her Erasmus internship. We also extend our appreciation to Dr. Michael Schell (Cenibra GmbH) for his support with the CQ1 microscope. Special thanks to OMNI Life Science and in particular to Dr. Markus Uhrig, for his generous support and provision of the CERO 3D Incubator and Bioreactor. We also thank Dr. Maik Wolfram-Schauerte for his critical feedback. This work was supported by the Imaging Assay Jumpstarter Contest Grant from Cenibra GmbH (awarded to AW and VA), a PhD scholarship from the Friedrich-Ebert-Stiftung and fellowships from the Joachim Herz Foundation and Bayer Foundation (awarded to AW). Research in the lab from LS is supported by institutional funds of the M3 Research Center and grants within the Cluster of Excellence (EXC2180), Image-guided and Functionally Instructed Tumor Therapies’ (iFIT) and the HECTOR Foundation MINT Personal Fonds.

## Author contributions

AW, VA and LS conceptualized the study. AW, VA and AM performed the experiments. VA implemented the aNSC culture, developed and *assembled* model and performed all light sheet microscopy. AW and VA jointly developed the *cancer core* models. AW developed the neurotoxicity screening assay and *advanced core*, as well as FACS protocols. AM supported cell culture maintenance and contributed to core model development. AW, VA and AM analyzed the data and interpreted the results. LS acquired major funding. AW and VA wrote the first draft of the manuscript. AW and VA jointly iterated and refined the text. CRN and LS supervised the project, provided critical feedback and corrected the manuscript. All authors reviewed and approved the final manuscript.

