## Supplemental Information for "3R-Compliant Murine *Ex Vivo* Platform for Scalable Brain Cancer Modeling and Drug Screening"

#### Contents

**1 General ... 3**

1.1 List of abbreviations ... 3

1.2 List of figures ... 4

1.3 List of videos ... 4

1.4 List of tables ... 4

**2 Materials and methods ... 5**

2.1 Materials ... 5

2.1.1 Consumables ... 5
2.1.2 Chemical compounds ... 7
2.1.3 Antibodies ... 7
2.1.4 Kits ... 7
2.1.5 Devices ... 8
2.1.6 Software ... 8

2.2 Methods ... 9

2.2.1 Harvesting murine adult neural stem cells (aNSCs) ... 9
2.2.2 Culture of adult neurospheres ... 9
2.2.3 Immunofluorescence staining and confocal microscopy ... 11
2.2.4 aNSCs differentiation into neurons, astrocytes and oligodendrocytes ... 11
2.2.5 Neurotoxicity assay in undifferentiated and differentiated aNSCs ... 12
2.2.6 Cultivation and maintenance of murine cell lines ... 13
2.2.7 2D cancer cell proliferation assay ... 13
2.2.8 Cultivation of cerebral organoids from aNSCs ... 14
2.2.9 Immunohistochemistry ... 15
2.2.10 Assembled model ... 16
2.2.11 Time-lapse live cell imaging via confocal microscopy ... 16
2.2.12 Light sheet microscopy ... 17
2.2.13 Cancer core model ... 18
2.2.14 Advanced cancer core model ... 20
2.2.15 Fluorescence-activated cell sorting (FACS) of cancer core models ... 21

**3 Supplementary figures ... 22**

Supplementary Figure S1 ... 22

Supplementary Figure S2 ... 23

Supplementary Figure S3 ... 25

Supplementary Figure S4 ... 27

Supplementary Figure S5 ... 29

Supplementary Figure S6 ... 30

Supplementary Figure S7 ... 31

**4 Supplementary videos ... 32**

**References ... 32**

### 1 General

#### 1.1 List of abbreviations

| | |
|---|---|
| 99LN | Breast-to-brain metastasis cell line 99LN BrM2 IVP (80478/79)[1] |
| ACC | Advanced cancer core |
| aNSC | Adult neural stem cells |
| AUF | Auranofin® |
| B-27 | B-27 neuronal cell culture supplement |
| B-27-A | B-27 neuronal cell culture supplement without vitamin A |
| BFP | Blue fluorescent protein |
| BSA | Bovine serum albumin |
| CC | Cancer core |
| DAPI | 4′,6-diamidino-2′-phenylindole |
| DBE | Dibenzyl ether |
| DMEM | Dulbecco's Modified Eagle Medium |
| DMSO | Dimethylsulfoxide |
| DPBS | Dulbecco's phosphate-buffered saline |
| EGF | Epidermal growth factor |
| FBS | Fetal bovine serum |
| FGF | Fibroblast growth factor |
| GFP | Green fluorescent protein |
| GL261 | Murine glioma cell line (Creative Biolabs, NCL-2108P28) |
| GlutaMAX | L-alanyl-L-glutamine dipeptide |
| H&E | Hematoxylin and eosin staining for the structure visualization |
| HBSS | Hank's balanced salt solution |
| KI-67 | Kiel 67 |
| IL-34 | Interleukin 34 |
| LLC1 | Lewis lung carcinoma LLC (ATCC, CRL-1642) |
| MK | Brain metastasis cells of melanoma cells K1735[2] |
| MTS | 3-(4,5-Dimethylthiazol-2-yl)-5-(3-carboxymethoxyphenyl)-2-(4-sulfophenyl)-2H-tetrazolium |
| Pen/Strep | Penicillin-Streptomycin |
| PFA | Paraformaldehyde |
| RFP | Red fluorescent protein |
| SVZ | Subventricular zone of the brain |
| TGF-β | Transforming growth factor β |
| TMZ | Temozolomide |
| ULA | Ultra-low attachment |

### 2 Materials and methods

#### 2.1 Materials

##### 2.1.1 Consumables

**Supplementary Table 1:** Consumables.

| Consumable | Supplier | Product number |
|---|---|---|
| 24-well ULA flat bottom plate | Corning | 3473 |
| 5 mL polypropylene tube | Greiner | 622201 |
| 5 mL polystyrene round-bottom tube with cell-strainer cap | Falcon | 352235 |
| 6-well suspension culture plate | Greiner | 657185 |
| 96-well clear flat bottom polystyrene TC-treated plate | Corning | 3596 |
| 96-well Nunclon Sphera-treated, U-shaped-bottom microplate | Nunclon | 174929 |
| Advanced DMEM/F12 | Gibco | 12634010 |
| Agarose | Promega | V2111 |
| Amber vial | Discountvials | CT242626-A-TEF-144 |
| Anti-adherence rinsing solution | STEMCELL Technologies | 07010 |
| Antibiotic antimycotic solution (100X), stabilized | Sigma-Aldrich | A5955 |
| B-27™ supplement (50X) | Gibco | 17504044 |
| B-27™ supplement (50X), minus vitamin A | Gibco | 12587010 |
| Bovine serum albumin (IgG-Free, protease-free) | Jackson ImmunoResearch | 001-000-162 |
| Cell culture dish, PS, 100/20 mm | Greiner | 664160 |
| Cell scraper | Corning | 3010 |
| Cellpath CellSafe+ biopsy capsule | Thermo Fisher Scientific Inc. | 15158886 |
| CERO tube | OMNI Life Science GmbH & Co KG | 2800005 |
| Counting chamber | Hycor | 87144 |
| Cryo tubes 2 mL | Greiner | 126263 |
| Cryomold | Weckert Labortechnik | 4565 |
| DAPI | Sigma-Aldrich | D9542 |
| Dispenser tips (1.25 mL) | Ratiolab | 29101102 |
| Dispenser tips (5 mL) | Ratiolab | 2910104 |
| Disposable graduated transfer pipets | VWR | 6124-543 |

| | | |
|---|---|---|
| Disposable scalpels | VWR | HERE1110810 |
| DMEM (1X) | Gibco | 21969-035 |
| DMEM (1X) without phenol red | Gibco | 31053-028 |
| DNA LoBind® tubes | Eppendorf | 0030108051 |
| Donkey serum | Jackson ImmunoResearch | 017-000-121 |
| DPBS (10X) | Gibco | 14200-067 |
| DPBS (1X) | Gibco | 14190-144 |
| Epredia™ Cassette II slotted tissue cassettes | Fisher Scientific | 16344146 |
| Falcon tubes 15 mL | Greiner Bio-One | 188271 |
| Falcon tubes 50 mL | Greiner | 227261 |
| Fetal bovine serum (FBS) Premium | PAN-Biotech | P30-3302 |
| Filter tips 10 µL | Greiner | 771353 |
| Filter tips 1000 µL | FisherBrand | 11973466 |
| Filter tips 20 µL | FisherBrand | 11943466 |
| Filter tips 200 µL | Greiner | 775353 |
| Filtercap cell culture flasks CELLSTAR® 175 $cm^2$ | Greiner | 660175 |
| Filtercap cell culture flasks CELLSTAR® 25 $cm^2$ | Greiner | 690175 |
| Filtercap cell culture flasks CELLSTAR® 75 $cm^2$ | Greiner | 658175 |
| GlutaMAX (35050061, Gibco) supplement | Gibco | 35050061 |
| HBSS | Gibco | 14025-126 |
| HEPES | Sigma-Aldrich | H3375 |
| Laminin | Sigma-Aldrich | L2020 |
| L-glutamine (200 mM/ 100X) | Gibco | 25030-024 |
| MACS SmartStrainer 100 µm | Miltenyi Biotec | 130-098-463 |
| mIL-34 | PeproTech | 315-34 |
| Mr. Frosty™ | Thermo Scientific™ | 5100-0001 |
| Neural stem cell basal medium | EMD Millipore | SCM003 |
| Penicillin-Streptomycin (5.000 U/mL) | Gibco | 15070063 |
| Pipets 10 mL | Greiner | 607180 |
| Pipets 25 mL | Greiner | 760180 |
| Pipets 5 mL | Greiner | 606180 |
| Poly-L-ornithine | Sigma-Aldrich | P4957 |
| Recombinant human FGF basic | PeproTech | 100-18B |
| Recombinant murine EGF | PeproTech | 315-09 |
| TGF-β | PeproTech | 315-01 |

| | | |
|---|---|---|
| Tissue culture plates 60 mm | Greiner | 628160 |
| Triton™ X-100 | Sigma-Aldrich | X-100 |
| TRIzol™ reagent | Invitrogen | 15596026 |
| Trypan blue | Carl Roth | CN76.3 |
| Trypsin inhibitor | Sigma-Aldrich | T6522 |
| Trypsin-EDTA (0.05%) | Gibco | 25300054 |
| Trypsin-EDTA (0.25%) | Gibco | 25200056 |

##### 2.1.2 Chemical compounds

**Supplementary Table 2:** Chemical compounds.

| **Compound** | **CAS** | **Supplier** | **Product number** | **Charge/ Batch** |
|---|---|---|---|---|
| Auranofin | 34031-32-8 | Sigma-Aldrich | A6733 | 0000144221 |
| Dibenzyl ether (DBE) | 103-50-4 | Sigma-Aldrich | 108014 | - |
| Dichloromethane | 75-09-2 | Sigma-Aldrich | 270997 | - |
| DMSO | 67-68-5 | Sigma-Aldrich | D2438 | - |
| Hydrogen peroxide | 7722-84-1 | Sigma-Aldrich | 216763 | - |
| Methanol | 67-56-1 | Fisher Chemical | A412SK-4 | - |
| ROTI®Histofix (4% Formal-dehyde) | 50-00-0 | Carl Roth | P087.3 | - |
| Temozolomide | 85622-93-1 | Tokyo Chemical Industry (TCI) Co., Ltd. | T2744 | - |
| Tween-20 | 9005-64-5 | Sigma-Aldrich | P9416 | - |

##### 2.1.3 Antibodies

**Supplementary Table 3:** Antibodies.

| **Antibody** | **Host-animal** | **Supplier** | **Product number** |
|---|---|---|---|
| Anti-GFAP | Goat | Abcam | ab53554 |
| Anti-MAP2 | Chicken | Synaptic Systems | 188 006 |
| Anti-Nestin | Rat | Sigma-Aldrich | MAB353 |
| Anti-NeuN | Rabbit | Abcam | ab177487 |
| Anti-Olig2 | Rabbit | Abcam | ab109186 |
| Anti-TUBB3 | Chicken | Abcam | ab41489 |
| Anti-SOX2 | Goat | R&D Systems | AF2018 |

##### 2.1.4 Kits

**Supplementary Table 4:** Kits.

| **Kit** | **Supplier** | **Product number** |
|---|---|---|
| CellTiter 96® AQueous One Solution Cell Proliferation Assay (MTS) | Promega | G3581 |

##### 2.1.5 Devices

**Supplementary Table 5:** Devices.

| Device | Supplier | Product number |
|---|---|---|
| CERO 3D Incubator & Bioreactor | Cell Microsystems | 2800000 |
| $CO_2$ resistant shaker (with 19 mm orbit) | Thermo Fisher Scientific Inc. | 88881101 |
| Vibratome Leica vt1200s | Leica Microsystems Vertrieb GmbH | 1491200S001 |
| X-Clarity Tissue Clearing System (XC- ETC1) | Logos Biosystems | C20001; C30001 |

##### 2.1.6 Software

**Supplementary Table 6:** Software.

| Software | Version |
|---|---|
| BioRender (www.biorender.com) | 2025 |
| CQ1 software | R1.08.02 |
| FlowJo | 10.10 |
| GraphPad | 10.1.0 |
| Microsoft Office Excel | 16.80 |
| Microsoft Office Word | 16.80 |

#### 2.2 Methods

##### 2.2.1 Harvesting murine adult neural stem cells (aNSCs)

The following procedure for harvesting aNSCs from murine brain tissue was adapted from Ahmed *et al.*:[3]

**Preparation and tissue isolation:**
6-24-week old mice were euthanized under the internal animal tissue collection reference number at the Institute for Tumor Biology and Experimental Therapy, Georg-Speyer-Haus, Frankfurt, Germany. Subsequently, the mouse heads, dissection area, and surgical tools were sterilized with 70% ethanol. The brain was excised and immediately placed in ice-cold dissection buffer consisting of HBSS (Gibco, 14025-126) and 2 mM HEPES (Sigma-Aldrich, H3375). In a laminar flow hood, the dentate gyrus (DG) region of hippocampus and subventricular zone (SVZ) were isolated, dissected, minced and suspended in dissection buffer on ice.

**Centrifugation and digestion:** The tissue was centrifuged at 200 g, 24°C for 1 minute. The supernatant was discarded and 1 mL of 0.05% trypsin-EDTA (Gibco, 25300054) was added. The sample was incubated at 37°C with gentle shaking for 15-20 minutes. Digestion was monitored and extended as necessary, ensuring not to exceed 30 minutes. After digestion, 1 mL of trypsin inhibitor (Sigma-Aldrich, T6522) was added, followed by an additional 20-minute incubation. Trituration was performed carefully using p1000 and p200 pipettes until a homogenous suspension was achieved.

**Washing:** The sample was diluted with 8 mL of wash media (Advanced DMEM/F12 (Gibco, 12634010), 1% Pen/Strep (Gibco, 15070063), 1% GlutaMAX (Gibco, 35050061)), then centrifuged at 200 g, 24°C for 5 minutes. The cells were then washed twice with 5 mL of wash media, followed by centrifugation under the same settings after each wash.

**Straining and seeding:** After the final wash, cells were resuspended in 6 mL of aNSC growth media (Neural stem cell basal medium (EMD Millipore, SCM003), 1% B27-A (Gibco, 12587010), 1% Antibiotic-Antimycotic (Sigma-Aldrich, A5955), 1% GlutaMAX (Gibco, 35050061), 7.5 µL EGF (100 µg/mL; PeproTech, 315-09), 7.5 µL FGF (100 µg/mL; PeproTech, 100-18B) per 25 mL media). The suspension was filtered through a 100 µm cell strainer (Miltenyi Biotec, 130-098-463) and centrifuged again. The resultant cell pellet was resuspended in 1 mL of aNSC growth media and seeded into a single well of a 24-well ultra-low attachment (ULA) culture plate (Corning, 3473).

**Incubation and media changes:** Cells were incubated at 37°C in a humidified 5% $CO_2$ incubator. Half of the media was replaced every other day, avoiding disruption of the neurospheres.

##### 2.2.2 Culture of adult neurospheres

This section describes two methods for culturing and passaging adult neurospheres (aNSCs), ensuring their maintenance at 50-100 µm in diameter and regular passaging every 2-4 days. The first method utilizes traditional 24-well ULA plates (Corning, 3473) or 25 cm² flasks (Greiner, 690175), while the second offers an alternative using the CERO 3D Incubator & Bioreactor (OMNI Life Science).

**Traditional culture method**

**Collection and enzymatic dissociation:** Neurospheres (50-100 µm in diameter) were collected using a p1000 pipette and transferred into a 5 mL tube. Cells were centrifuged at 200 g, 24°C for 5 minutes. The supernatant was aspirated and 1 mL of 0.05% trypsin-EDTA (Gibco, 25300054) was added to each tube and incubated at 37°C with slow mixing for 2 minutes. An equal volume of trypsin inhibitor (Sigma-Aldrich, T6522) was added and the cells were triturated first with a p1000 pipette, then with a p200 for finer dissociation.

**Centrifugation and resuspension:** Following trituration, the cell mixture was centrifuged again at 200 g, 24°C for 5 minutes. The supernatant was aspirated and the cell pellet was resuspended in aNSC growth media (Neural stem cell basal medium (EMD Millipore, SCM003), 1% B27-A (Gibco, 12587010), 1% Antibiotic-Antimycotic (Sigma-Aldrich, A5955),1% GlutaMAX (Gibco, 35050061), 7.5 µL EGF (100 µg/mL; PeproTech, 315-09), 7.5 µL FGF (100 µg/mL; PeproTech, 100-18B) per 25 mL media).

**Seeding:** Finally, the cells were seeded into a 24-well ULA culture plate (Corning, 3473) at 1 mL per well or into a 25 $cm^2$ culture flask (Greiner, 690175) with 3.5-5 mL of aNSCs growth media (same composition as above) per flask. Cells were incubated at 37°C in a 5% $CO_2$ humidified incubator, with passaging every 2-4 days.

**Alternative method using CERO 3D Incubator & Bioreactor**

The CERO 3D Incubator & Bioreactor provides a more efficient alternative, allowing larger-scale cultures with the capacity to grow up to 50 million aNSCs per tube in 10 mL of media.

**Procedure:** The cells were passaged following the same initial steps as above to generate a single-cell suspension. After resuspension, the suspension was transferred into a CERO tube (OMNI Life Science, 2800005) with at least 12 mL of aNSCs growth media (same composition as above).

The CERO Bioreactor was set to rotate 1 second clockwise and 1 second counterclockwise with continuous motion, providing optimal mixing and preventing cell aggregation during incubation.

**Advantages of CERO bioreactor:**

- **Efficient growth:** The continuous rotation of the CERO system keeps cells in suspension afloat, preventing premature clumping and enabling rapid expansion, with several million cells forming per tube.
- **Neurosphere formation:** After 3-5 days, well-formed neurospheres will develop. Passaging is recommended once neurospheres reach 50-100 µm in diameter, approximately every week.
- **Optional cerebral organoid formation:** Larger neurospheres can serve as starting material for cerebral organoid development, streamlining workflows for organoid culture.

##### 2.2.3 Immunofluorescence staining and confocal microscopy

Immunofluorescence (IF) and imaging of sectioned organoid samples involves the following steps:

**Dewaxing:** Organoid sections on glass slides were dewaxed by the core facility to remove any paraffin residues.

**Hydrophobic circle drawing:** Once dried, a hydrophobic barrier was created around each specimen using a pap pen liquid blocker.

**Blocking:** Samples were blocked using a blocking solution containing 3% BSA (001-000-162, Jackson ImmunoResearch) and 0.01% Triton-X (Sigma-Aldrich, X-100) at 4°C for 1 hour to prevent non-specific binding of antibodies.

**Antibody incubation:** Post-blocking, slides were dried and the primary antibody mix (diluted 1:1000 in a 1:1 PBS/Blocking solution) was applied. The slides were then incubated overnight within a wet chamber placed in the fridge. After primary antibody incubation, slides were dried and washed three times with PBS for 5 minutes. The slides were dried and then covered with the secondary antibody mix (approximately 200 µL) for 1 hour in a wet chamber The secondary antibody solution was prepared at a 1:500 dilution in a 1:1 PBS/blocking solution.

**DAPI staining:** After secondary antibody incubation, slides were dried and then washed twice with 1X PBS (Gibco, 14190-144). DAPI (D9542, Sigma-Aldrich) was applied at a 1:5000 dilution in a 1:1 PBS/Blocking solution for 5 minutes to stain the nuclei, followed by three PBS washes.

**Mounting and storage**: The slides were carefully dried and a drop of mounting media was added directly onto the specimen before covering with a coverslip. The samples were left to dry overnight in a semi-open, light-protected dry chamber at room temperature. The slides were stored in the fridge in the dark to preserve the fluorescence signal until imaging.

**Imaging:** Prepared slides were imaged using the CellVoyager CQ1 (Yokogawa) in suitable magnification for respective application.

##### 2.2.4 aNSCs differentiation into neurons, astrocytes and oligodendrocytes

aNSCs were differentiated into neurons, astrocytes and oligodendrocytes as follows:

**Preparation of culture surface:** 96-well plates (TC-treated, flat bottom) (Corning, 3596) were coated with 20 µg/mL poly-L-ornithine solution (Sigma-Aldrich, P4957) in 1X PBS (Gibco, 14190-144) and incubated at 37°C for at least 30 minutes. Plates were washed three times with 1X PBS. Subsequently, 5 µg/mL laminin solution (Sigma-Aldrich, L2020) in 1X PBS was added and the plate was incubated at 37°C for at least 30 minutes. A final 1X PBS wash was performed to prepare the wells for cell seeding.

**Cell seeding:** aNSCs were harvested and dissociated into single cells as per the previously described method (see **Section 2.2.2**). Cells were seeded using M1 media (Adv. DMEM/F12

(Gibco, 12634010), 1% B27 (Gibco, 17504044), 1% L-Gln (Gibco, 25030-024), 1% GlutaMAX (Gibco, 35050061), 1% Pen/Strep (Gibco, 15070063), 7.5 µL EGF (100 µg/mL; PeproTech, 315-09), 7.5 µL FGF (100 µg/mL; PeproTech, 100-18B) per 25 mL media) into the poly-L-ornithine/laminin-coated wells. The seeded cells were maintained at 37°C in a humidified environment containing 5% $CO_2$.

**Differentiation process:** Starting the day after seeding, half of the media was replaced daily with freshly prepared differentiation media. The differentiation media consisted of Adv. DMEM/F12, 1% B27, 1% Antibiotic-Antimycotic, 1% GlutaMAX, 2.5 µL EGF (100 µg/mL), 2.5 µL FGF (100 µg/mL) per 25 mL media.

This differentiation process was continued for 3-5 days, allowing for the differentiation of neurons, astrocytes and oligodendrocytes.

##### 2.2.5 Neurotoxicity assay in undifferentiated and differentiated aNSCs

**Plate coating:** 96-well plates (Corning, 3596) were coated with 20 µg/mL poly-L-ornithine solution (Sigma-Aldrich, P4957) and incubated at 37°C for at least 30 minutes. The coated wells were washed three times with 1X PBS (Gibco, 14190-144) for 1 minute each. Subsequently, wells were coated with 5 µg/mL laminin solution (Sigma-Aldrich, L2020) at 37°C for 30 minutes, followed by a final 1X PBS wash before cell seeding.

**Cell seeding:** Single-cell suspensions of aNSCs (50-100 µm) were prepared as described in **Section 2.2.2**. Cell counts were determined using a Neubauer chamber (Hycor, 87144) after brief staining with Trypan Blue (Carl Roth, CN76.3). For undifferentiated aNSCs, $6 \times 10^3$ cells were seeded per well in 200 µL of aNSCs growth media (Neural stem cell basal medium (EMD Millipore, SCM003), 1% B27-A (Gibco, 12587010), 1% Antibiotic-Antimycotic (Sigma-Aldrich, A5955),1% GlutaMAX (Gibco, 35050061), 7.5 µL EGF (100 µg/mL; PeproTech, 315-09), 7.5 µL FGF (100 µg/mL; PeproTech, 100-18B) per 25 mL media) and incubated overnight before treatment. For differentiated aNSCs, $6 \times 10^3$ cells were seeded in 200 µL of M3 media (Adv. DMEM/F12 (Gibco, 12634010), 1% B27 (Gibco, 17504044), 1% Pen/Strep (Gibco, 15070063), 1% GlutaMAX (Gibco, 35050061), 2.5 µL EGF (100 µg/mL; PeproTech, 315-09), 2.5 µL FGF (100 µg/mL; PeproTech, 100-18B) per 25 mL media) and allowed to grow for 3-5 days before treatment.

**Treatment phase (first 48 hours):** Cells were treated with temozolomide (TMZ, 500 µM) or Auranofin® (AUF, 2 µM) for 48 hours, with media-only and vehicle-only (1% DMSO, v/v) controls. Fresh media containing the respective treatments was added and plates were incubated at 37°C for 3, 6, 12, 24 and 48 hours. 2 hours prior to each time point, 20 µL of MTS reagent (Promega, G3581) was added per well and incubated at 37°C on an orbital shaker (Thermo Fisher, 88881101) in a humidified 5% $CO_2$ incubator. The absorbance was measured at 490 nm and 630 nm using a microplate reader.

**Recovery phase (48 + 24 h and 48 + 48 h):** At the 48-hour mark, drug-containing media was completely replaced with fresh media to simulate drug clearance. Plates were incubated for an additional 24 or 48 hours, with viability assessed at 48+24 h and 48+48 h using the MTS assay.

All results were normalized to media-only controls of the respective timepoint (set to 100%), with background absorbance ($A_{630nm}$) subtracted from formazan production ($A_{490nm}$).

#### 2.2.6 Cultivation and maintenance of murine cell lines

Murine cell lines were cultured using the following standard cell culture procedures.

**Thawing procedure:** Frozen cell lines were rapidly thawed at 37°C, then transferred to a 15 mL Falcon tube (Greiner, 188271) containing 9 mL of pre-warmed adv. DMEM compl. media (Adv. DMEM/F12 (Gibco, 12634010), 10% FBS (PAN-Biotech, P30-3302), 1% L-Gln (Gibco, 25030-024), 1% Pen/Strep (Gibco, 15070063)). After centrifugation at 1500 rpm, room temperature for 3 minutes, the supernatant was aspirated and the cell pellet was resuspended in fresh adv. DMEM compl. media (same composition as above) before being seeded into a T25 flask (Greiner, 690175).

**Culture maintenance:** Cells were incubated at 37°C in a humidified incubator with 5% $CO_2$. Media was refreshed every third day and cells were passaged when reaching 80% confluency to maintain nutrient levels and cell health. Confluent cells were passaged by removing the media, washing with 1X PBS (Gibco, 14190-144) and incubating the cells with 0.05% trypsin-EDTA (Gibco, 25300054) for 2-5 minutes at 37°C. The reaction was neutralized with DMEM compl. media (DMEM (Gibco, 21969-035), 10% FBS (PAN-Biotech, P30-3302), 1% L-Gln (Gibco, 25030-024), 1% Pen/Strep (Gibco, 15070063)) and the cells were centrifuged at 1500 rpm, 4°C for 3 minutes. The cell pellet was resuspended in fresh media, with part of the suspension transferred to a new flask for continued culture and the rest reserved for experimental assays.

**Cryopreservation:** For long-term storage, cells were split as per the passaging procedure. The final cell pellet was resuspended in adv. DMEM compl. media (same composition as above) supplemented with 10% DMSO (Sigma-Aldrich, 67-68-5), aliquoted into cryovials (Greiner, 126263) and gradually cooled to -80°C using a Mr.Frosty™ (Thermo Scientific™, 5100-0001) before transfer to liquid nitrogen for a longer storage.

#### 2.2.7 2D cancer cell proliferation assay

**Cell preparation and seeding:** Murine glioblastoma (GL261) cells were detached using 0.05% trypsin-EDTA (Gibco, 25200056), neutralized with DMEM compl. media (DMEM (Gibco, 21969-035), 10% FBS (PAN Biotech, P30-3302), 1% L-Gln (Gibco, 25030-024), 1% Pen/Strep (Gibco, 15070063)) and counted using a Neubauer chamber (Hycor, 87144) after Trypan Blue staining (Carl Roth, CN76.3). A total of $1 \times 10^4$ cells per well were seeded in a 96-well plate (Corning, 3596) containing phenol red-free DMEM compl. (DMEM (Gibco, 31053-028), 10% FBS, 1% L-Gln, 1% Pen/Strep) and incubated at 37°C in a humidified 5% $CO_2$ incubator overnight to allow adhesion.

**Treatment phase (first 48 hours):** Cells were treated with temozolomide (TMZ, 500 µM) or Auranofin® (AUF, 2 µM) for 48 hours, with media-only and vehicle-only (1% DMSO, v/v) controls. Fresh media containing the respective treatments was added and plates were incubated at 37°C for 3, 6, 12, 24 and 48 hours. 2 hours prior to each time point, 20 µL of MTS reagent (Promega, G3581) was added per well and incubated at 37°C on an orbital shaker (Thermo Fisher, 88881101) in a humidified 5% $CO_2$ incubator. The absorbance was measured at 490 nm and 630 nm using a microplate reader.

**Recovery phase (48 + 24 h and 48 + 48 h):** At the 48-hour mark, drug-containing media was completely replaced with fresh media to simulate drug clearance. Plates were incubated for an additional 24 or 48 hours, with viability assessed at 48+24 h and 48+48 h using the MTS assay.

All results were normalized to media-only controls of the respective timepoint (set to 100%), with background absorbance ($A_{630nm}$) subtracted from formazan production ($A_{490nm}$).

##### 2.2.8 Cultivation of cerebral organoids from aNSCs

Cerebral organoids were generated from aNSCs using two approaches: an orbital shaker method and the CERO 3D Incubator & Bioreactor (OMNI Life Science) for enhanced efficiency and scalability.

**Method 1: Cultivation using an orbital shaker**
**Neurosphere splitting and cell quantification:** Neurospheres were dissociated into single-cell suspensions following the procedure outlined in **Section 2.2.2**. Cell numbers were determined using a Neubauer chamber (Hycor, 87144) after staining with Trypan Blue (Carl Roth, CN76.3).

**Organoid seeding:** aNSCs were seeded at a density of $5x10^3$-$2x10^4$ cells per well in a U-bottom, 96-well ULA plate (Nunclon, 174929) in 200 µL of M1 media (Adv. DMEM/F12 (Gibco, 12634010), 1% B27 (Gibco, 17504044), 1% L-Gln (Gibco, 25030-024), 1% GlutaMAX (Gibco, 35050061), 1% Pen/Strep (Gibco, 15070063), 7.5 µL EGF (100 µg/mL; PeproTech, 315-09), 7.5 µL FGF (100 µg/mL; PeproTech, 100-18B) per 25 mL media).

**Medium replacement:** After 72 hours, 100 µL of M1 media was aspirated and replaced with M2 media (Adv. DMEM/F12, 1% B27, 1% L-Gln, 1% GlutaMAX, 1% Pen/Strep, 5 µL EGF (100 µg/mL), 5 µL FGF (100 µg/mL) per 25 mL media). After another 72 hours, 100 µL of M2 media was replaced with M3 media (Adv. DMEM/F12, 1% B27, 1% Pen/Strep, 1% GlutaMAX, 2.5 µL EGF (100 µg/mL), 2.5 µL FGF (100 µg/mL) per 25 mL media).

**Organoid transfer and maintenance:** After 72 hours in M3 media, organoids were transferred to 6-well suspension plates (Greiner, 657185) with up to 3 mL of M3 media (same composition as above) per well. Media changes were conducted every 72 hours for continued growth and maintenance.

**Method 2: Cultivation using the CERO 3D Incubator & Bioreactor**
The CERO 3D Incubator & Bioreactor offers an alternative to traditional methods, promoting round, uniform organoid growth with reduced hands-on time and increased scalability. The larger culture volume also minimizes the frequency of media changes.

**Pre-seeding in ULA plates:** A single-cell suspension from neurospheres was prepared as per the procedure in **Section 3.2**. Cell counts were determined using a Neubauer chamber after Trypan Blue staining. $5x10^3$-$2x10^4$ cells were seeded per well in a 96-well ULA plate (Nunclon, 174929) in 200 µL of M1 media (same composition as above).

**Transfer to CERO tubes:** After 3 days, organoids were transferred from ULA plates to CERO tubes (OMNI Life Science, 2800005) with 12 mL M2 media (same composition as above) or pre-incubated in 6-well suspension plates (Greiner, 657185) with up to 3 mL M2 media per well before transfer to CERO tubes.

**Media changes and incubation:** Media changes were performed twice per week, replacing half of the media or adding 3 mL of fresh media if the medium retains a healthy color. After 1 week, the media was switched to M3 media (same composition as above) for all subsequent media changes. Organoids grew beyond 2 mm in diameter within 3 months, with long-term cultivation up to 5 months being possible.

**Incubation parameters:**

- For initial growth, set the CERO Bioreactor to rotate 1 second clockwise and 1 second counterclockwise continuously to ensure proper mixing.
- Once organoids reach 250 µm in diameter, adjust the rotation to 2 seconds clockwise, pause for 1 second, then 2 seconds counterclockwise, to maintain optimal culture conditions as organoids grow larger.

**Alternative method: Direct neurosphere outgrowth into organoid culture**
An alternative approach for simplifying organoid generation involves allowing aNSCs to transition directly from neurospheres to organoids.

**Procedure:** Once neurospheres had reached 100 µm in diameter, the neurosphere media was replaced with M1 media (same composition as above). After 1 week, during the subsequent media change, the media was switched to M2 media (same composition as above). After another week, M3 media (same composition as above) was used for all subsequent changes.

This method reduces hands-on time and materials while improving reproducibility.

##### 2.2.9 Immunohistochemistry

**Sample fixation and processing:** Culture media was aspirated from the sample wells and samples were washed 1-2 times with 1X PBS (Gibco, 14190 144). Fixation was performed by fully covering the samples with ROTI®Histofix (4% formaldehyde) (Carl Roth, P087.3) and incubating them at room temperature for 24-48 hours. Following fixation, samples were carefully transferred using a single-use transfer pipette into CellSafe+ Biopsy capsules (Fisher Scientific Inc., 15158886). Capsules were then placed into Epredia™ Cassette II slotted tissue cassettes (Fisher Scientific, 16344146), securely closed and submerged in 70% ethanol (EtOH) for preservation.

**Tissue processing and staining:** Samples were sent to the HistoCore Facility at the M3 Research Center, Tübingen, where they underwent dehydration, paraffin embedding and sectioning into 2-3 µm slices. Sections were then deparaffinized and stained with Hematoxylin and Eosin (H&E) for morphological assessment. Immunohistochemistry (IHC) staining was performed using Ki67, a marker for cellular proliferation and Cleaved Caspase-3 (CC3) to assess apoptosis. Stained sections were imaged using a brightfield digital pathology scanner (e.g., Leica Aperio AT2).

##### 2.2.10 Assembled model

To generate assembled models, pre-differentiated cerebral organoids (1 month) were fused with pre-formed cancer spheroids (3 days), combining healthy brain tissue with pathological spheroids. The detailed protocols are outlined below.

**Spheroid culture:** Spheroids were generated from a single cancer cell suspension (see **Section 2.2.6**). After cell counting with a Neubauer chamber (Hycor, 87144), cells were centrifuged at 1500 rpm, room temperature for 5 minutes and pellet was resuspended in spheroid-specific media (DMEM compl. or DMEM (Gibco, 21969-035), 1% L-Gln (Gibco, 25030-024), 1% Pen/Strep (Gibco, 15070063) and optionally 12.5 ng/mL EGF (100 µg/mL; PeproTech, 315-09)). Cells were seeded into a U-bottom, 96-well ULA plate (Nunclon, 174929) at 200 µL medium per well. The plate was incubated at 37°C in a humidified 5% $CO_2$ CO2 incubator for 72 hours.

**Assembled model formation (organoid-spheroid co-culture):** For assembled model formation, pre-formed cerebral organoids (Actb-DsRed or WT) of comparable size were placed into a 96-well ULA plate (Nunclon, 174929). Pre-formed cancer spheroids were introduced into each well containing an organoid. The co-cultures were incubated under the same conditions as the spheroids (37°C, 5% $CO_2$) for 72 hours to allow assembled model formation.

**Live imaging:** The development of assembled models was monitored in real time using benchtop fluorescence microscope with imaging performed every 24 hours to track progression.

##### 2.2.11 Time-lapse live cell imaging via confocal microscopy

Real-time live cell imaging was performed to monitor organoids or assembloids over time using confocal microscopy. This method ensured optimal environmental conditions while minimizing sample disruption during long-term imaging.

**Preparation of organoid/ assembloid samples:** Individual organoids or assembloids were prepared for imaging in a U-bottom, 96-well ULA plate (Nunclon, 174929). For each well, a single organoid or assembloid was placed and covered with 200 µL of M3 media (Adv. DMEM/F12 (Gibco, 12634010), 1% B27 (Gibco,17504044), 1% Pen/Strep (Gibco, 15070063), 1% GlutaMAX (Gibco, 35050061), 2.5 µL EGF (100 µg/mL; PeproTech, 315-09), 2.5 µL FGF (100 µg/mL; PeproTech, 100-18B)). To reduce media evaporation during prolonged imaging sessions, all unused wells were filled with sterile water or PBS.

**Microscope setup:** The prepared 96-well plate was transferred to a confocal microscope equipped with integrated environmental control systems, including temperature regulation (37°C), $CO_2$ monitoring (5% $CO_2$) and humidity control to ensure sample viability. Imaging parameters were defined based on experimental requirements: wells containing organoids or assembloids were selected for imaging, fluorescence channels were assigned according to molecular reporters (e.g., GFP, RFP) and time-lapse acquisition was configured for the desired duration (e.g., 24-72 hours) with intervals between captures (e.g., every 60 minutes). To balance imaging detail with phototoxicity, appropriate magnification, resolution and Z-stack settings were applied where 3D imaging was required.

Automated time-lapse imaging protocols were generated using the microscope software, enabling multi-channel fluorescence acquisition and Z-stack capture as necessary. Throughout the experiment, environmental conditions (temperature, $CO_2$ and humidity) were continuously monitored to maintain sample stability. The imaging session was initiated and periodic checks were performed to ensure stable operation and proper data acquisition.

**Post-imaging workflow:** Acquired images were saved at regular intervals and exported to a secure storage location for analysis. Image processing and quantitative analysis were performed using software such as ImageJ or Imaris. Key metrics, including organoid growth, structural changes and fluorescence intensity, were extracted for further evaluation.

##### 2.2.12 Light sheet microscopy

This protocol, adapted from a previously published method,[4] is optimized for imaging whole organoids and / or co-culture systems using light sheet microscopy.

**Collection of organoids:** Organoids were harvested using a 3 mL transfer pipette and transferred into amber glass vials (Discountvials, CT242626-A-TEF-144) for processing.

**Pretreatment of organoid samples:** Organoids were dehydrated in a graded methanol series (20%, 40%, 60%, 80%, 100%; Fisher Chemical, A412SK-4), then incubated in 100% methanol at 4°C. Clearing was achieved by incubation in 66% dichloromethane (Sigma-Aldrich, 270997) and 33% methanol overnight, followed by bleaching in 5% $H_2O_2$ (Sigma-Aldrich, 216763) in methanol. Rehydration through a reverse methanol series (1 hour per step) was followed by two washes in PTx.2 buffer[4] at room temperature to prepare for immunostaining.

**Immunolabeling of pretreated organoids:** Samples were permeabilized in permeabilization buffer[4] at 37°C for 2 days, then blocked with a solution of 42 mL PTx.2 buffer, 3 mL Donkey serum (Jackson Immunoresearch, 017-000-121) and 5 mL DMSO (Sigma-Aldrich, 67-68-5) at 37°C for 2 more days. Organoids were incubated with primary antibodies in PTwH (PBS (Gibco, 14190-144), 0.2% Tween-20 (Sigma-Aldrich, P9416), 5% DMSO (Sigma-Aldrich, D2438)) at 37°C for 2 days. After washing 4-5 times in PTwH, samples were incubated with secondary antibodies in PTwH with 3% Donkey serum (Jackson ImmunoResearch, 017-000-121) at 37°C for 2 days. Excess secondary antibodies were removed by washing 4-5 times in PTwH.

**Clearing of organoids:** Samples were embedded in 1.5% low-melting agarose (Promega, V2111) for stabilization, then dehydrated through a graded methanol series (20%, 40%, 60%, 80%, 100%; Fisher Chemical, A412SK-4), with each step lasting 1 hour (or optionally overnight). Clearing was performed with 66% dichloromethane (Sigma-Aldrich, 270997) and 33% methanol for 3 hours, followed by two 15-minute washes in 100% dichloromethane (Sigma-Aldrich, 270997). Samples were finally incubated in dibenzyl ether (DBE) (Sigma-Aldrich, 108014), ensuring the tube was filled almost completely to prevent oxidation. Before imaging, the tube was gently inverted to mix the solution. Organoids were then ready for light sheet microscopy.

##### 2.2.13 Cancer core model

This method outlines the establishment of the cancer core model, which integrates cancer cells with differentiated cerebral organoids. It provides a reproducible system with precise control over cell ratios and tumor cell growth, effectively modeling distinct tumor microenvironments (TMEs).

**Tested cell ratios:**
The following cell ratios were tested to study how varying cancer cell ratios influence core formation (**Supplementary Table 7)**:

**Supplementary Table 7:** Cancer core model seeding ratios.

| Cerebral organoids (cells/well) | Cancer cells (cells/well) | Cancer core ratio | Cancer core formation |
|---|---|---|---|
| $2x10^4$ | $1x10^4$ | 2:1 | Mosaic-like pattern with cancer cells and non-cancerous cells throughout the whole model |
| $4x10^4$ | $1x10^4$ | 4:1 | Cancer core unit with high cancer cell density |
| $4x10^4$ | $2x10^4$ | 2:1 | Mosaic-like pattern with cancer cells and non-cancerous cells throughout the whole model |
| $8x10^4$ | $2x10^4$ | 4:1 | Cancer core unit with high cancer cell density |
| $1x10^5$ | $1x10^3$ | 100:1 | Late-onset patch-like pattern with cancer cells distributed throughout the cerebral organoid structure |
| $1x10^5$ | $5x10^3$ | 20:1 | Patch-like pattern with cancer cells distributed throughout the cerebral organoid structure |
| $1x10^5$ | $1x10^4$ | 10:1 | Cancer core unit with high cancer cell density with larger cerebral 'shell' |
| $1x10^5$ | $2.5x10^4$ | 4:1 | Cancer core unit with high cancer cell density |

These tested cell ratios led to distinct cancer core model patterns (healthy cerebral cells vs. pathological cancer cells):

- 2:1 ratio → Produces a mosaic-like pattern with cancer cells distributed throughout the structure and no well-defined cerebral 'shell'.
- 4:1 ratio → Forms a core unit pattern with a clearly defined cerebral 'shell'.
- 10:1 ratio → Generates a dense cancer core unit with a high concentration of cancer cells and a larger cerebral 'shell'.
- 20:1 ratio → Creates a patch-like pattern where cancerous cell form "patches" within the cerebral organoid structure.

- 100:1 ratio → Produces a late-onset patch-like pattern with cancer cells distributed throughout the cerebral organoid structure.

**Influence of cancer cell type:**
The cancer core formation also strongly depends on the biological background of the cancer cell line:

- Glioma model (GL261): Exhibits a highly infiltrative pattern, with cancer cells spreading widely throughout the cerebral organoid, mimicking the invasive nature of gliomas observed clinically.
- Breast-to-brain metastasis model (99LN): Tends to form well-defined, localized cancer regions within the cerebral organoid, reflecting the distinct tumor morphology seen in breast-to-brain metastases.
- Lung-to-brain metastasis model (LLC1): Highly invasive, with fast-growing cancer areas that resemble metastatic cancer spread in the brain.
- Melanoma-to-brain metastasis model (MK1735): Tends to form more localized and well-defined cancer areas within the organoid structure.

**Cancer core formation:** Cancer cells (e.g., GL261-GFP, 99LN-BFP, LLC1-GFP or MK-BFP) were prepared from 2D culture following standard passaging procedures (see **Section 2.2.6**). Cells were counted using a Neubauer chamber (Hycor, 87144) and viability was assessed with Trypan Blue (Carl Roth, CN76.3). The cancer cells were resuspended in M3 media (Adv. DMEM/F12 (Gibco, 12634010), 1% B27 (Gibco, 17504044), 1% Pen/Strep (Gibco, 15070063), 1% GlutaMAX (Gibco, 35050061), 2.5 µL EGF (100 µg/mL; PeproTech, 315-09), 2.5 µL FGF (100 µg/mL; PeproTech, 100-18B) per 25 mL media). The resulting suspension was filtered through a 100 µm strainer (Miltenyi Biotec, 130-098-463) to ensure uniformity.

**Organoid dissociation:** In parallel, pre-differentiated cerebral organoids (grown for a minimum of 1 month) were enzymatically dissociated to create a single-cell suspension to integrate with cancer cells. In detail, fifty to one hundred cerebral organoids (diameter > 300 µm) were transferred into a 15 mL Falcon tube and centrifuged at 200 g for 2 minutes. The media was aspirated and 1-2 mL of 0.05% trypsin-EDTA (Gibco, 25300054) was added. The organoid suspension was incubated at 37°C on an orbital shaker (Thermo Fisher, 88881101) for 2 minutes. After incubation, the organoids were gently triturated with a p1000 pipette, followed by a p200 pipette, until a uniform single-cell suspension was achieved. To stop the enzymatic reaction, 1 mL of trypsin inhibitor (Sigma-Aldrich, T6522) was added. The suspension was centrifuged again at 200 g for 2 minutes and the pellet was resuspended in 1-2 mL of M3 media (same composition as above). The resulting suspension was filtered through a 100 µm strainer (Miltenyi Biotec, 130-098-463) to ensure uniformity.

**Final cancer core model assembly:** Depending on the desired cerebral organoid-to-cancer cell ratio, mixtures were prepared using ratios ranging from 2:1 to 100:1 (cerebral organoid cells to cancer cells). These mixtures were distributed into a 96-well ULA plate (Nunclon, 174929) at 200 µL per well. The plates were incubated at 37°C in a humidified 5% $CO_2$ incubator on an orbital shaker for 72 hours. The cancer core model was monitored using benchtop fluorescence microscope to track the interaction and integration of cancer cells with organoid-derived cells.

##### 2.2.14 Advanced cancer core model

This method outlines the development of advanced cancer core (ACC) model, which integrates tumor cells, dissociated cerebral organoid cells and immune cells - particularly microglia - within a cerebral organoid structure. This model more accurately represents TME, enabling a detailed study of tumor-immune interactions and cancer progression.

**Tested cell ratios:** The following cell ratios were tested, all leading to defined model formation (**Supplementary Table 8**). Unlike the cancer core, the advanced cancer core initially shows a mosaic-like pattern at 24 hours, with the cerebral shell closing over the next 48 hours to form the advanced cancer core.

**Supplementary Table 8:** ACC model seeding ratios.

| Cerebral organoids (cells/well) | Cancer cells (cells/well) | Microglia cells (cells/well) | ACC Ratio |
|---|---|---|---|
| $2x10^4$ | $1x10^4$ | $1x10^4$ | 2:1:1 |
| $4x10^4$ | $1x10^4$ | $2x10^4$ | 4:1:2 |
| $4x10^4$ | $2x10^4$ | $2x10^4$ | 2:1:1 |
| $8x10^4$ | $2x10^4$ | $4x10^4$ | 4:1:2 |

**Influence of cancer cell type:** The advanced cancer core formation depends highly on the biological background of the cancer cell line:

- Glioma model (GL261): Exhibits a highly infiltrative pattern with substantial microglia presence within the tumor. This mimics the clinical observation where approximately 30% tumor mass contain microglia.
- Breast-to-brain metastasis model (99LN): The advanced cancer core exhibits less defined tumor borders compared to the standard 99LN cancer core. As seen in clinical samples, microglia are found only at the tumor periphery, leading to peripheral microgliosis rather than direct tumor infiltration.

**Advanced cancer core formation**: Cancer cells (e.g., GL261-GFP, 99LN-BFP) were cultured using the standard passaging technique (see **Section 2.2.6**). Microglia (EOC2 cells) were cultured separately in DMEM compl. media (DMEM (Gibco, 21969-035), 10% FBS (PAN-Biotech, P30-3302), 1% L-Gln (Gibco, 25030-024), 1% Pen/Strep (Gibco, 15070063)) supplemented with 50 µL Interleukin-34 (mIL-34) (100 µg/mL; PeproTech, 315-34) and 2.5 µL TGF-β (50 µg/mL; PeproTech, 315-01) per 25 mL media to maintain cell viability.

**Simultaneous preparation of single-cell suspensions:** Cerebral organoids (diameter > 300 µm, grown for at least 1 month) were transferred into a 15 mL Falcon tube and centrifuged at 200 g for 2 minutes. The supernatant was aspirated and the organoids were resuspended in 1-2 mL of 0.05% trypsin-EDTA (Gibco, 25300054). They were incubated at 37°C on an orbital shaker for 2 minutes. Organoids were then gently triturated using a p1000 pipette, followed by a p200 pipette, until a uniform single-cell suspension was achieved. The reaction was stopped by adding trypsin inhibitor (Sigma-Aldrich, T6522) and the suspension was centrifuged at 200 g for 2 minutes. The pellet was resuspended in modified M3 media (Adv. DMEM/F12 (Gibco, 12634010), 1% B27 (Gibco,17504044), 1% Pen/Strep (Gibco, 15070063), 1% GlutaMAX (Gibco, 35050061), 2.5 µL EGF (100 µg/mL; PeproTech, 315-09), 2.5 µL FGF (100 µg/mL; PeproTech, 100-18B), 50 µL IL-34 (100 µg/mL; PeproTech, 315-34), 2.5 µL TGF-β (50 µg/mL; PeproTech, 315-01) per 25 mL media). Cancer cells were detached using 0.05% trypsin-EDTA (Gibco, 25300054), while microglia (EOC2 cells) were detached using a cell scraper (Corning, 3010). Both were resuspended in modified M3 media (same composition as above). All cell types were counted using Neubauer chambers (Hycor, 87144) and their viability was assessed using Trypan Blue (Carl Roth, CN76.3) exclusion.

**Mastermix preparation:** A mastermix was prepared according to the desired cell ratios (e.g., 2:1:1, 4:1:2) of cerebral organoid cells, cancer cells and microglia. The mastermix was added into a U-bottom, 96-well ULA plate (Nunclon, 174929) at 200 µL per well.

**Seeding and incubation:** After seeding, the plates were immediately placed on an orbital shaker inside a 37°C incubator with 5% $CO_2$. The plates were shaken at 100 rpm for 72 hours to promote advanced cancer core formation.

**Monitoring and clinical relevance:** The advanced cancer core model was monitored using benchtop fluorescence microscopy to observe the integration of immune cells, tumor cells and neuronal cells.

##### 2.2.15 Fluorescence-activated cell sorting (FACS) of cancer core models

**Initial preparation of samples:** Six to nine cancer core models per experimental condition were pooled and centrifuged at 200 g, 4°C for 5 minutes to pellet the cells. The supernatant was discarded and the cell pellet was resuspended in 1-1.5 mL of 0.25% trypsin-EDTA (Gibco, 25200056) for dissociation. The cells were incubated on an orbital shaker (Thermo Fisher, 88881101) at 37°C in a humidified 5% $CO_2$ incubator for 5 minutes. Trituration was then performed to achieve a single-cell suspension.

**Cell suspension processing:** Trypsin inhibitor (Sigma-Aldrich, T6522) was added. The suspension was centrifuged and the supernatant discarded. The pellet was resuspended in 500 µL of 1X PBS (Gibco, 14190-144). The suspension was passed through a 100 µm strainer (Milteniy Biotec, 130-098-463), ensuring a uniform single-cell suspension for FACS.

**FACS:** For each sample, 1.5 mL DNA LoBind tubes (Eppendorf, 0030108051) were prepared with 50 µL of 1X PBS (Gibco, 14190-144). To exclude dead cells, samples were stained with DAPI (Sigma-Aldrich, D9542) for 5 minutes before FACS. The gating strategy was designed to exclude DAPI-positive cells ($DAPI^+$; dead cells), enabling the sorting of two final populations: DAPI-negative/GFP-positive ($DAPI^-$/ $GFP^+$; live cancer cells) and DAPI-negative/RFP-positive ($DAPI^-$/ $RFP^+$; live organoid cells) using a BD FACSAria™ Fusion Flow Cytometer (see **Supplementary Figure S5 & S6**).

**Post-FACS processing:** A minimum of $5x10^4$ cells per sample were sorted and immediately kept on ice. TRIzol™ Reagent (Invitrogen, 15596026) was added for RNA preservation and samples were stored at -80°C for subsequent RNA isolation.

#### 3 Supplementary figures

##### Supplementary Figure S1

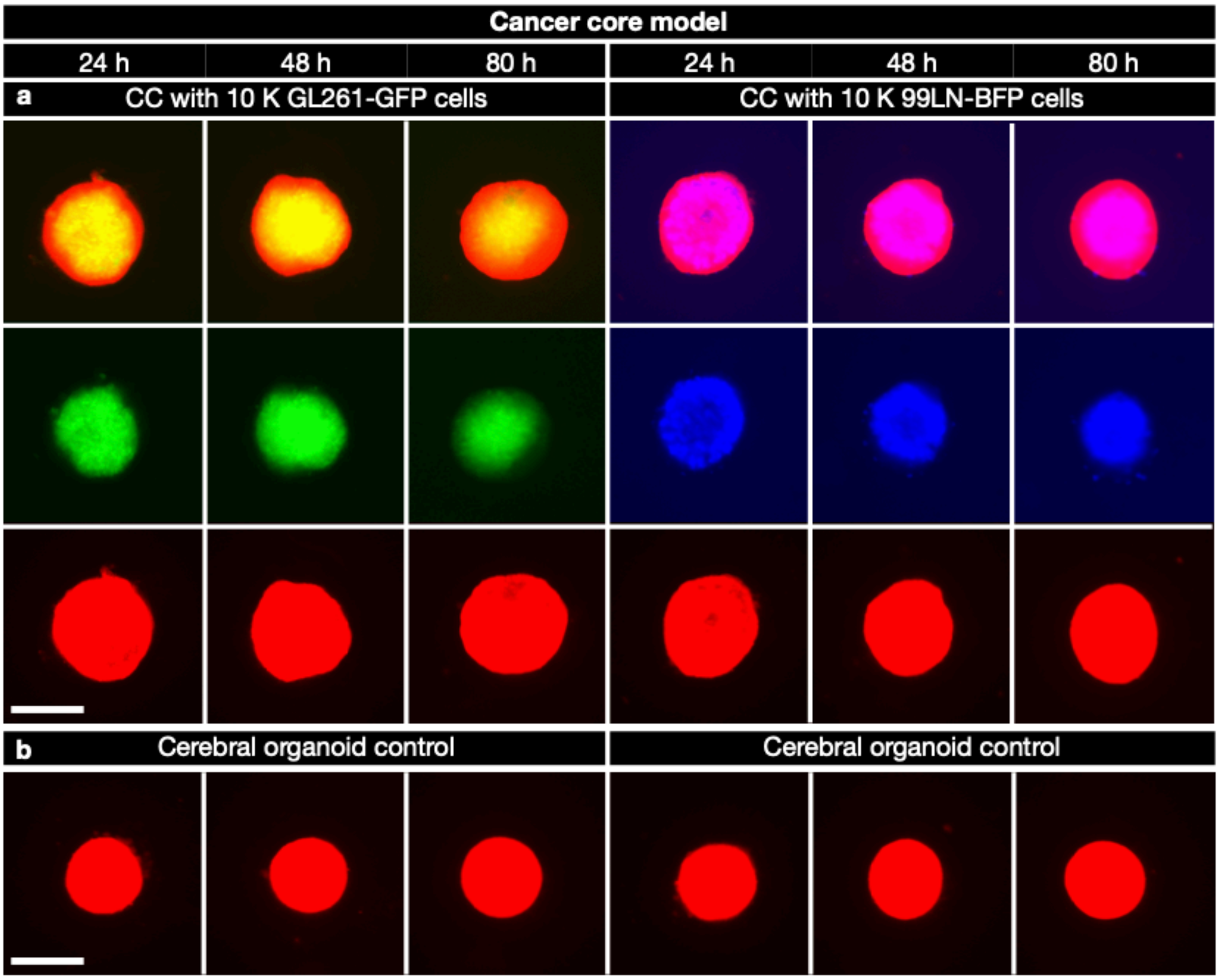

**Supplementary Figure S1:** Tumor core organization and cerebral organoid viability in cancer core models and controls. **a**, Cancer core models generated by co-culturing $1x10^5$ RFP-labeled cerebral organoid cells with $1x10^4$ GL261-GFP glioma cells (right) or $1x10^4$ 99LN-BFP breast-to-brain metastasis cells (left). Cancer cores exhibit well-defined tumor masses fully surrounded by cerebral organoid cells at 24, 48 and 80 hours. **b**, Control cerebral organoid models seeded with $1x10^5$ RFP-labeled cerebral organoid cells from a single-cell suspension. Within 24 hours, cells reform complex 3D architectures, demonstrating robust viability, with continued growth and structural maturation over 80 hours. Scale bar: 500 µm.

#### Supplementary Figure S2

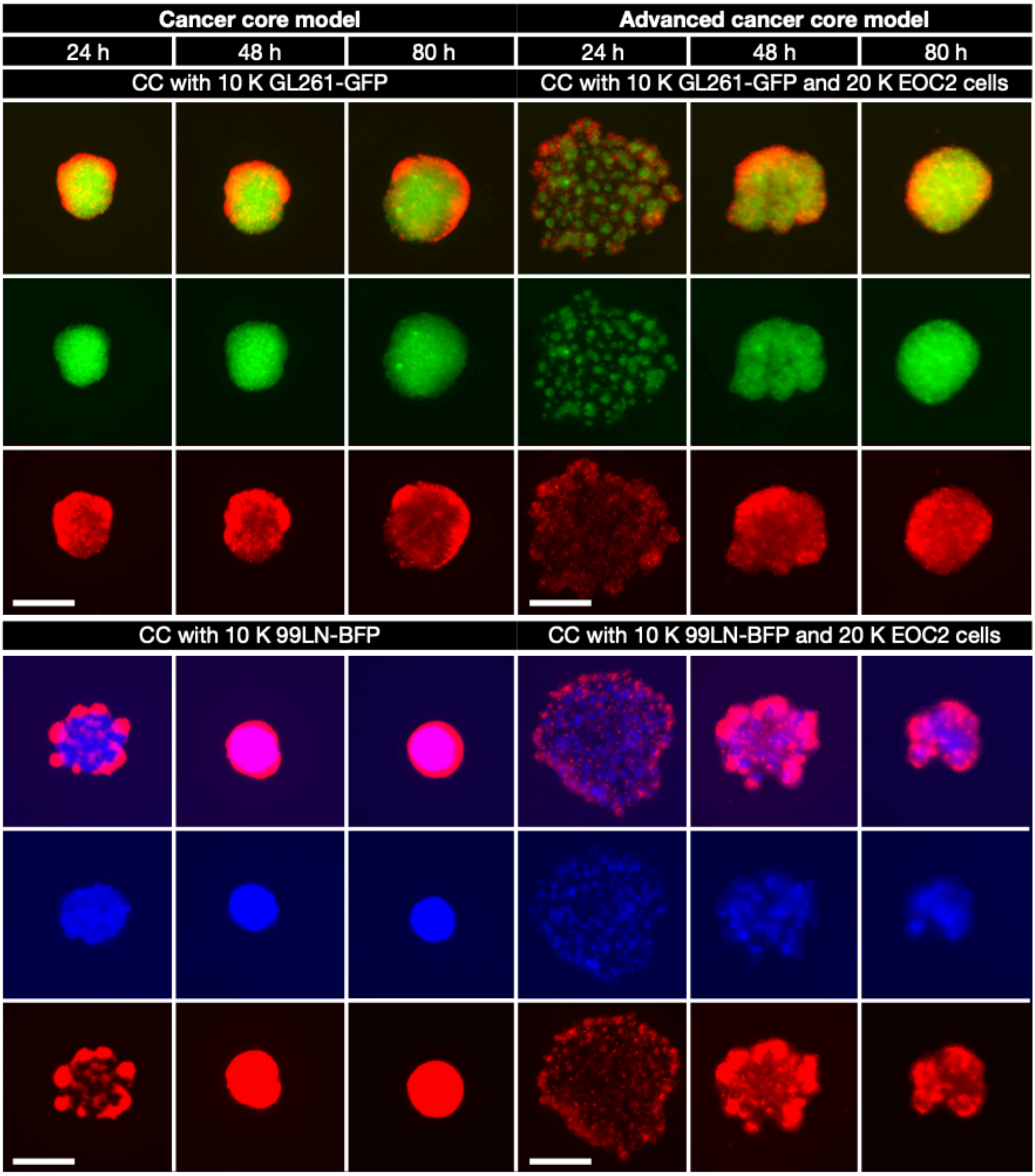

**Supplementary Figure S2:** Immune cell integration alters tissue organization in glioma and breast-to-brain metastasis CC models (4:1 and 4:1:2 ratios). **Top left:** Glioma CC without EOC2, generated by co-culturing $4 \times 10^4$ RFP-labeled cerebral organoid cells with $1 \times 10^4$ GL261-GFP tumor cells (4:1 ratio). **Top right:** Glioma ACC with EOC2 microglia, incorporating $4 \times 10^4$ RFP-labeled cerebral organoid cells, $1 \times 10^4$ GL261-GFP tumor cells and $2 \times 10^4$ EOC2 microglia (4:1:2 ratio). **Bottom left:** Breast-to-brain metastasis CC without EOC2, formed by co-culturing $4 \times 10^4$ RFP-labeled cerebral organoid cells with $1 \times 10^4$ 99LN-BFP tumor cells (4:1 ratio). **Bottom right:** Breast-to-brain metastasis ACC with EOC2 microglia, integrating $4 \times 10^4$ RFP-labeled cerebral organoid cells, $1 \times 10^4$ 99LN-BFP tumor cells and $2 \times 10^4$ EOC2 microglia (4:1:2 ratio). Scale bar: 500 μm. Images captured at 24, 48 and 80 hours.

#### Supplementary Figure S3

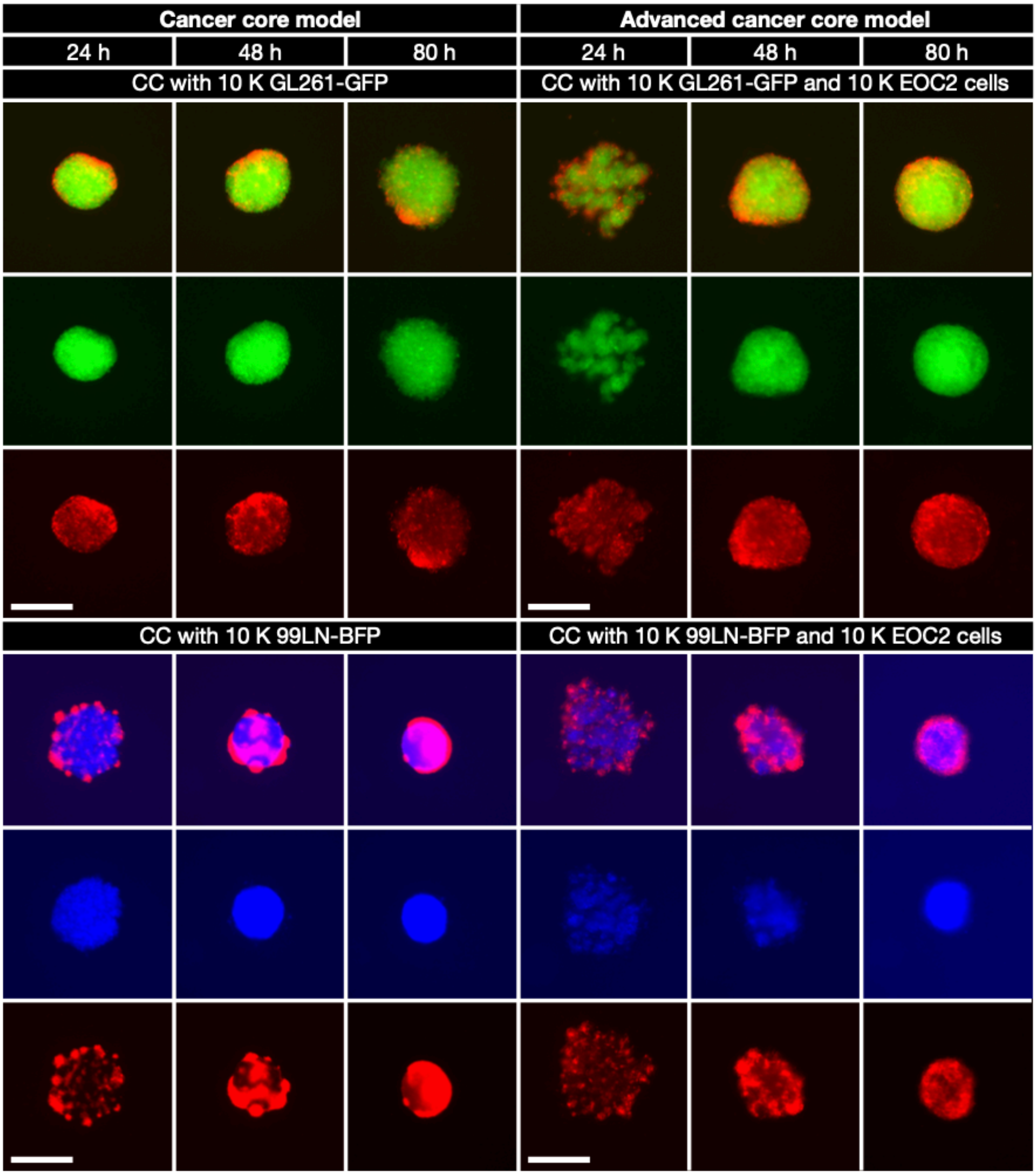

**Supplementary Figure S3:** Immune cell integration alters tissue organization in glioma and breast-to-brain metastasis cancer core (CC) models (2:1 and 2:1:1 ratios). **Top left:** Glioma CC without EOC2, generated by co-culturing 2x$10^4$ RFP-labeled cerebral organoid cells with 1x$10^4$ GL261-GFP tumor cells (2:1 ratio). **Top right:** Glioma ACC with EOC2 microglia, incorporating 2x$10^4$ RFP-labeled cerebral organoid cells, 1x$10^4$ GL261-GFP tumor cells and 1x$10^4$ EOC2 microglia (2:1:1 ratio). **Bottom left:** Breast-to-brain metastasis CC without EOC2, formed by co-culturing 2x$10^4$ RFP-labeled cerebral organoid cells with 1x$10^4$ 99LN-BFP tumor cells (2:1 ratio). **Bottom right:** Breast-to-brain metastasis ACC with EOC2 microglia, integrating 2x$10^4$ RFP-labeled cerebral organoid cells, 1x$10^4$ 99LN-BFP tumor cells and 1x$10^4$ EOC2 microglia (2:1:1 ratio). Scale bar: 500 µm. Images captured at 24, 48 and 80 hours.

#### Supplementary Figure S4

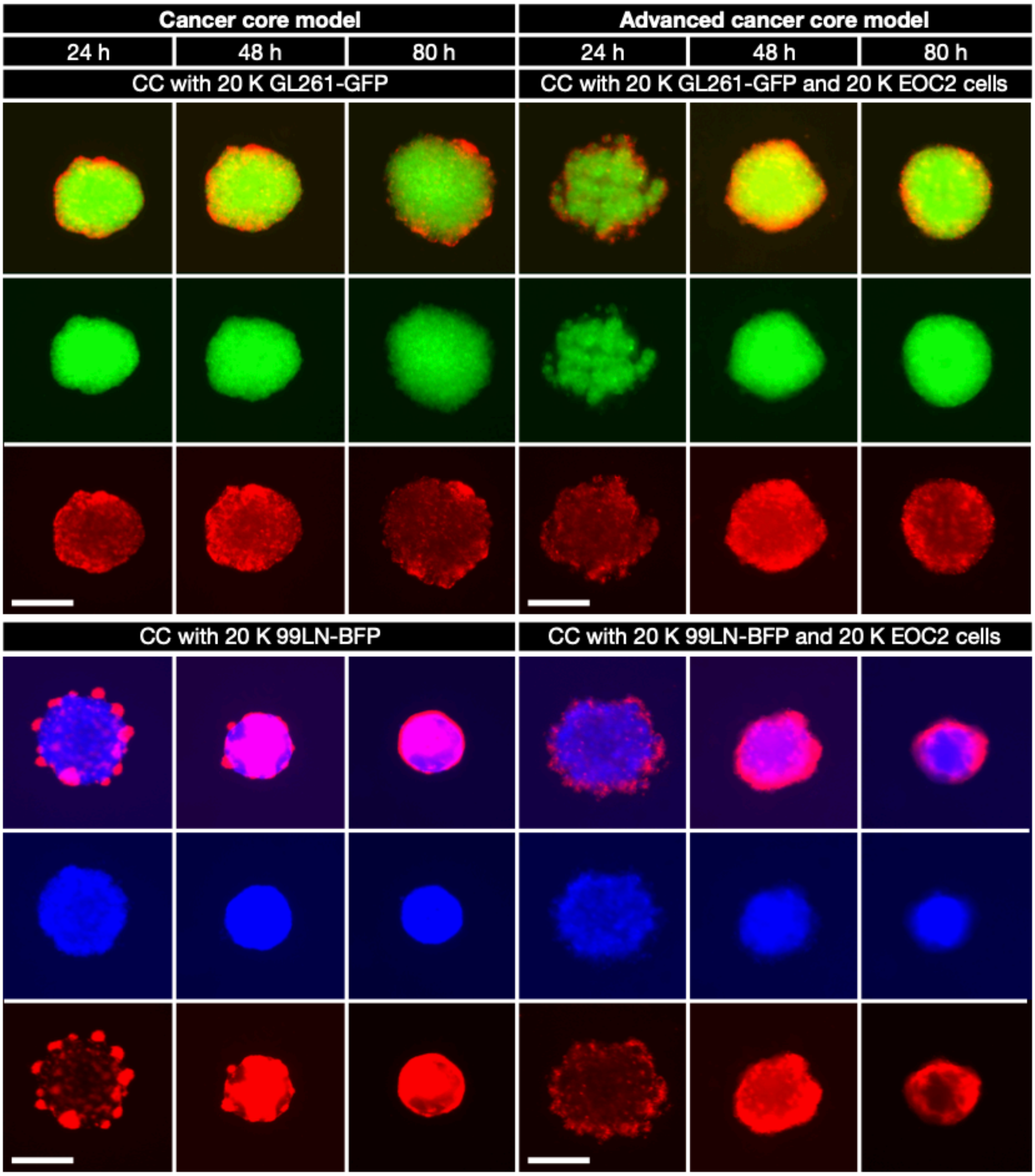

**Supplementary Figure S4:** Similar tissue organization alterations in CC models with higher cell seeding numbers after immune cell integration. **Top left:** Glioma CC without EOC2, generated by co-culturing $4x10^4$ RFP-labeled cerebral organoid cells with $2x10^4$ GL261-GFP tumor cells (2:1 ratio). **Top right:** Glioma ACC with EOC2 microglia, incorporating $4x10^4$ RFP-labeled cerebral organoid cells, $2x10^4$ GL261-GFP tumor cells and $2x10^4$ EOC2 microglia (2:1:1 ratio). **Bottom left:** Breast-to-brain metastasis CC without EOC2, formed by co-culturing $4x10^4$ RFP-labeled cerebral organoid cells with $2x10^4$ 99LN-BFP tumor cells (2:1 ratio). **Bottom right:** Breast-to-brain metastasis ACC with EOC2 microglia, integrating $4x10^4$ RFP-labeled cerebral organoid cells, $2x10^4$ 99LN-BFP tumor cells and $2x10^4$ EOC2 microglia (2:1:1 ratio). Scale bar: 500 µm. Images captured at 24, 48 and 80 hours.

#### Supplementary Figure S5

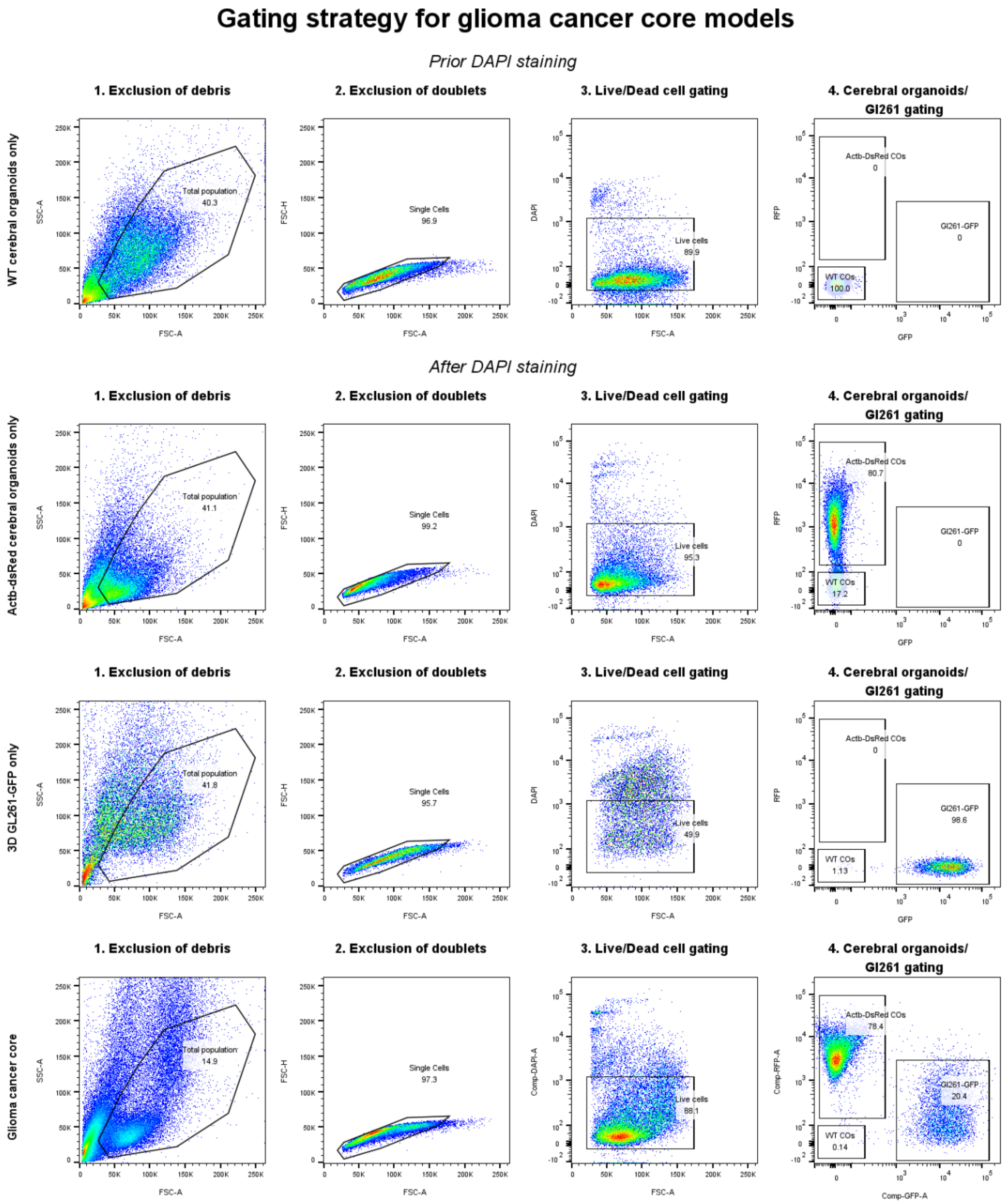

**Supplementary Figure S5:** Gating strategy for FACS-based isolation of viable cell populations in glioma cancer core models. Flow cytometry gating strategy for isolating viable cell populations. **Row 1:** Wild-type (WT) cerebral organoids prior to DAPI staining. **Row 2:** Actb-DsRed cerebral organoids showing RFP-positive populations with DAPI staining to identify live (DAPI-negative) cells. **Row 3:** GL261-GFP cells identifying GFP-positive cancer cells. **Row 4:** Glioma cancer core or breast-to-brain metastasis cancer core, enabling isolation of viable non-cancerous organoid cells (RFP-positive, DAPI-negative) and cancerous cells (GFP-positive, DAPI-negative).

#### Supplementary Figure S6

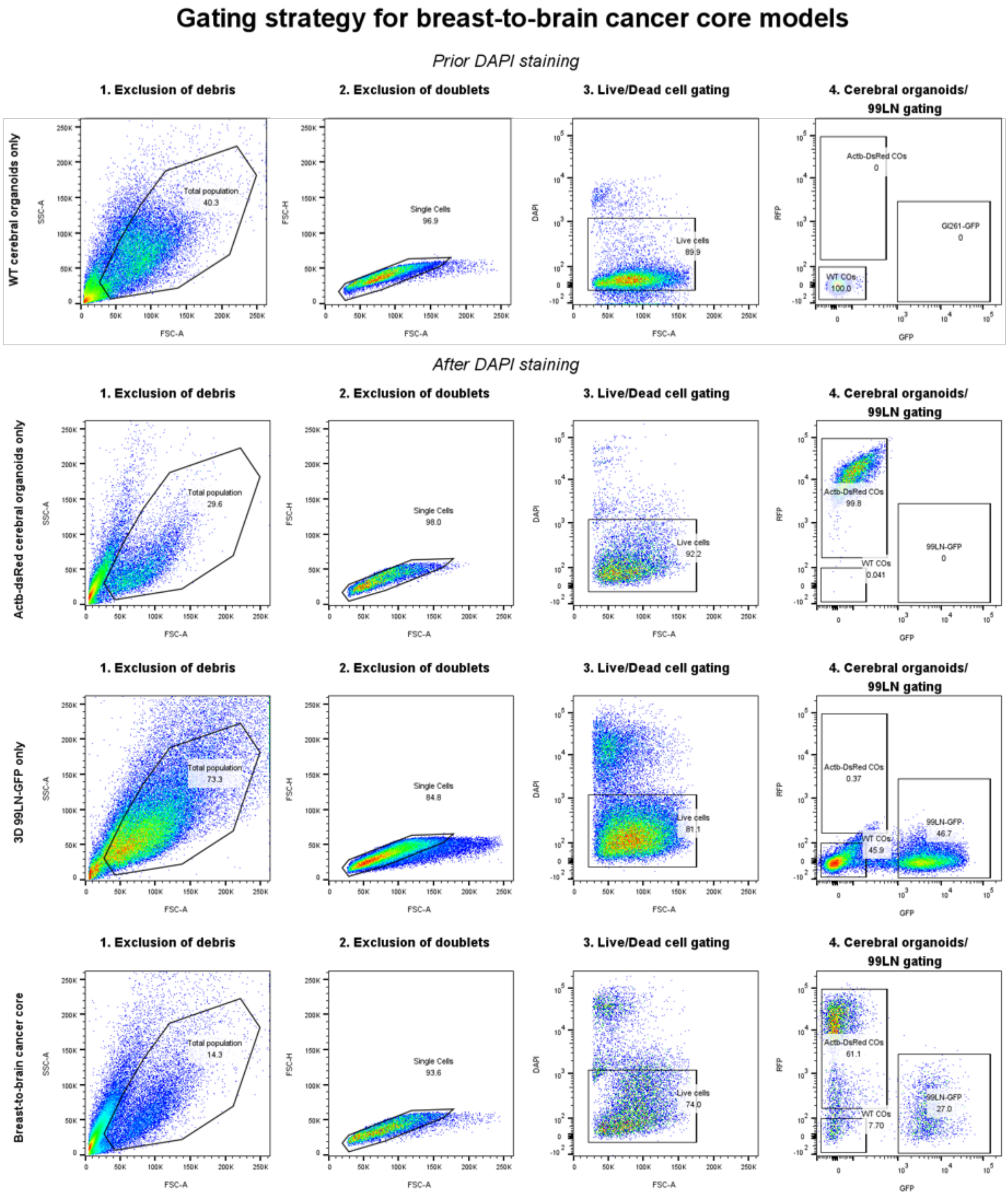

**Supplementary Figure S6:** Gating strategy for FACS-based isolation of viable cell populations in breast-to-brain metastasis cancer core models. Flow cytometry gating strategy for isolating viable cell populations. **Row 1:** Wild-type (WT) cerebral organoids prior to DAPI staining. **Row 2:** Actb-DsRed cerebral organoids showing RFP-positive populations with DAPI staining to identify live (DAPI-negative) cells. **Row 3:** 99LN-GFP cells identifying GFP-positive cancer cells. **Row 4:** Breast-to-brain metastasis cancer core, enabling isolation of viable non-cancerous organoid cells (RFP-positive, DAPI-negative) and cancerous cells (GFP-positive, DAPI-negative).

#### Supplementary Figure S7

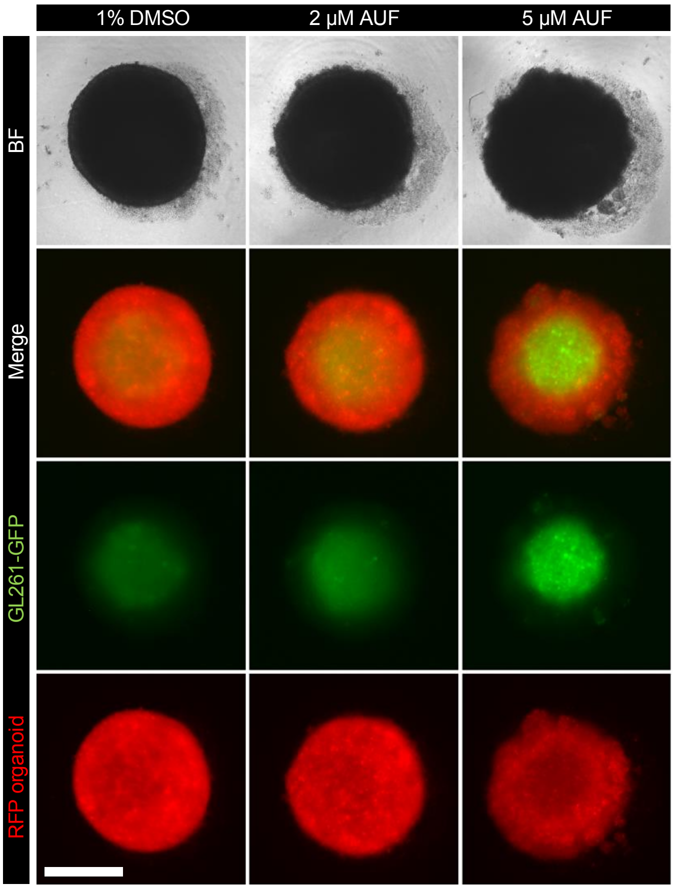

**Supplementary Figure S7:** Glioma cancer core model for combined drug efficacy and neurotoxicity screening. Glioma cancer core models were generated by co-culturing $1x10^5$ RFP-labeled cerebral organoid cells with $1x10^4$ GL261-GFP glioma cells and grown for 72 hours prior to treatment. Models were treated with either vehicle-only control (1% DMSO, v/v; left column), 2 μM Auranofin® (AUF; middle column), or 5 μM AUF (right column) and imaged 24 hours post-treatment. Images are shown in four rows: **Row 1:** Brightfield (BF), **Row 2:** Merged IF images of RFP-labeled cerebral organoid cells and GL261-GFP tumor cells, **Row 3:** GL261-GFP tumor cells (green), **Row 4:** RFP-labeled cerebral organoid cells (red). Compared to controls, AUF-treated cancer cores exhibit reduced rim integrity, with the 5 μM treatment showing the greatest decrease in the cerebral organoid shell. Scale bar: 500 μm.

#### 4 Supplementary videos

**Supplementary Video 1:** Time-lapse imaging of cerebral organoids co-cultured with GFP-labeled GL261 glioma spheroids. RFP-labeled cerebral organoids ($1x10^5$ cells; red) were co-cultured with GFP-labeled GL261 glioma spheroids ($1x10^4$ cells; yellow). Time-lapse imaging over 72 hours, captured at 2-hour intervals, showed extensive tumor-cell interactions, with glioma cells progressively invading the organoid. Scale bar: 200 µM.

**Supplementary Video 2:** Time-lapse imaging of organoids co-cultured with GFP-labeled LLC1 lung-to-brain metastatic spheroids. RFP-labeled cerebral organoids ($1x10^5$ cells; red) were co-cultured with GFP-labeled LLC metastasis spheroids ($1x10^4$ cells; turquoise). Time-lapse imaging over 72 hours, at 2-hour intervals, revealed rapid, distal invasion into the organoid, showing aggressive tumor dynamics and structural disruption of neural tissue. Scale bar: 200 µM.

**Supplementary Video 3:** Time-lapse imaging of cerebral organoids co-cultured with BFP-labeled MK melanoma-to-brain spheroids. RFP-labeled cerebral organoids ($1x10^5$ cells; red) were co-cultured with MK-BFP melanoma-to-brain spheroids ($1x10^4$ cells; blue). Imaging over 72 hours, at 2-hour intervals, captured minimal spheroid growth while cerebral cells moved around the spheroid. Scale bar: 200 µM.

**Supplementary Video 4:** Time-lapse imaging of cerebral organoids co-cultured with GFP-labeled 99LN breast-to-brain spheroids. RFP-labeled cerebral organoids ($1x10^5$ cells; red) were co-cultured with GFP-labeled 99LN breast-to-brain spheroids ($4x10^4$ cells; orange). Time-lapse imaging over 72 hours, with 2-hour intervals, revealed breast-to-brain cancer cells forming compact spheroids with delayed infiltration into the cerebral tissue. Scale bar: 200 µM.

**Supplementary Video 5:** Light-sheet imaging of a cancer core model. A cancer core model comprising RFP-labeled cerebral organoids ($1x10^5$ cells; red) and GFP-labeled 99LN breast-to-brain metastatic cells ($1x10^3$ cells; orange) was cultured for 10 days, fixed and cleared. Light-sheet imaging revealed the tumor infiltration into the organoid with a distinct patch-like distribution pattern, highlighting spatial tumor organization within the model. Scale bar: 200 µM.
